# DLL4 and PDGF-BB regulate migration of human iPSC-derived skeletal myogenic progenitors

**DOI:** 10.1101/2021.02.28.431778

**Authors:** Giulia Ferrari, SungWoo Choi, Louise A. Moyle, Kirsty Mackinlay, Naira Naouar, Christine Wells, Francesco Muntoni, Francesco Saverio Tedesco

**Author notes:** Equally contributing authors (SungWoo Choi, Louise A. Moyle). Institute of Biomedical Engineering, University of Toronto, Toronto, M5S3E1, Canada (Louise A. Moyle). University of Cambridge, Department of Physiology, Development and Neuroscience, Cambridge CB23DY, UK.

## Abstract

Muscle satellite stem cells (MuSCs) are responsible for skeletal muscle growth and regeneration. Despite their differentiation potential, human MuSCs have limited *in vitro* expansion and *in vivo* migration capacity, limiting their use in cell therapies for diseases affecting multiple skeletal muscle groups such as muscular dystrophies. Several protocols have been developed to derive progenitor cells similar to MuSCs from human induced pluripotent stem cells (hiPSCs), in order to establish a source of myogenic cells with controllable proliferation and differentiation capacity. However, currently available hiPSC myogenic derivatives also suffer from limitations of cell migration, ultimately delaying their clinical translation. Here we provide evidence that activation of NOTCH and PDGF pathways with DLL4 and PDGF-BB improves migration of hiPSC-derived myogenic progenitors *in vitro*. Transcriptomic and functional analyses demonstrate that this property is conserved across species and multiple hiPSC lines, including genetically-corrected hiPSC derivatives from a patient with Duchenne muscular dystrophy. DLL4 and PDGF-BB treatment had no negative impact on cell proliferation; cells maintained their myogenic memory, with differentiation fully rescued by NOTCH inhibition. RNAseq analysis indicate that pathways involved in cell migration are modulated in treated myogenic progenitors, consistent with results from functional profiling of cell motility at single cell resolution. Notably, treated cells also showed enhanced trans-endothelial migration in transwell assays. Enhancing extravasation is a key translational milestone for intravascular delivery of hiPSC myogenic derivatives: our study establishes the foundations of a transgene-free, developmentally inspired strategy to achieve this goal, moving hiPSCs one step closer to future muscle gene and cell therapies.

## INTRODUCTION

Muscle satellite stem cells (MuSCs) reside between the basal lamina and sarcolemma of muscle fibres and are responsible for growth and regeneration of skeletal myofibres. Upon activation, MuSCs give rise to an activated progeny named myoblasts, which then repair and regenerate myofibres (reviewed in (Benedetti, Hoshiya and Tedesco, 2013)). Myoblasts have been tested in numerous clinical trials for Duchenne muscular dystrophy (DMD), the most common muscular dystrophy of childhood, which severely affects most skeletal muscles and remains incurable (reviewed in (Tedesco *et al*., 2010)). However, despite promising pre-clinical results in animal models, to date, only myogenic cell therapies of localised muscular dystrophies such as oculopharyngeal muscular dystrophy (OPMD) have reported functional improvements upon myoblast transplantations in patients (Périé *et al*., 2014).

Skeletal myogenic cells have been delivered via the intramuscular or the intravascular route (Tedesco *et al*., 2010). However, the efficacy of both transplantation modalities is impeded by insufficient migration, leading to poor muscle biodistribution of donor cells. Intramuscular injections frequently result in generation of chimeric myofibres often limited to the area adjacent to the needle trajectory, necessitating multiple injections and making this strategy unfeasible for generalised myopathies such as DMD (Skuk, 2004). On the other hand, intravascular delivery of donor cells via major arteries may facilitate simultaneous targeting of multiple muscle groups. Intra-arterial injections of mesoangioblasts, myogenic cells derived from a subset of muscle perivascular cells, ameliorated muscle pathology and function in pre-clinical models of muscular dystrophy and was also translated into a phase I/IIa clinical trial in five DMD boys (Cossu *et al*., 2016). Although mesoangioblasts are still considered promising candidate cells for systemic delivery owing to their trans-endothelial migration (also known as extravasation) capacity, they possess lower skeletal myogenic and self-renewal capacity than MuSCs, which limits their long-term translational potential. Therefore, an ideal cell type for myogenic cell therapies should possess the migratory capacity of perivascular cells as well as the differentiation and self-renewing potential of MuSCs.

NOTCH signalling plays a pivotal role in cell fate specification during embryonic myogenesis, as well as in post-natal MuSC self-renewal and differentiation (Conboy and Rando, 2002; Schuster-Gossler, Cordes and Gossler, 2007; Bjornson *et al*., 2012; Mourikis and Tajbakhsh, 2014; Baghdadi *et al*., 2018; Verma *et al*., 2018). Canonical NOTCH signalling involves interactions between NOTCH ligands (e.g., Delta-like (DLL) 1, 3, 4 and Jagged (JAG) 1, 2) and receptors (NOTCH 1-4). Perturbation of NOTCH signalling in donor cells has shown context-dependent effects on myogenic cell transplants. Treatment of mouse and human myoblasts on culture dishes coated with DLL1 and DLL4 did not enhance engraftment in *mdx* mice, a DMD mouse model (Sakai *et al*., 2017). However, DLL1 treatment of canine MuSCs maintained their engraftment potential during *in vitro* expansion (Parker *et al*., 2012). Furthermore, modulation of the DLL1-NOTCH1 axis in both mouse and human mesoangioblasts supported improvement of the dystrophic phenotype after intra-arterial delivery in mice (Quattrocelli *et al*., 2014). Platelet-derived growth factor (PDGF) signalling is another important regulator of myogenic cell behaviour. PDGF receptor-β (PDGFR-β) is expressed by cells derived from the mesenchyme (Dellavalle *et al*., 2007; Trojanowska, 2009). PDGF-BB, the putative ligand of PDGFR-β, is expressed by endothelial cells and dystrophic muscle fibres for recruitment of pericytes and MuSCs, respectively (Betsholtz, 2004; Piñol-Jurado *et al*., 2017).

Previous work showed that mouse embryonic myoblasts in close proximity to blood vessels undergo a spontaneous fate shift into pericyte-like cells in vivo; this phenomenon was mimicked *in vitro* by treating embryonic myoblasts with DLL4 and PDGF-BB (Cappellari *et al*., 2013). More recently, we showed that modulation of NOTCH and PDGF pathways induces perivascular cell features while enhancing self-renewal and migration in adult mouse and human MuSC-derived myoblasts (Gerli *et al*., 2019). However, the translational potential of primary, tissue-derived MuSCs is hindered by the need to obtain them invasively (i.e. via muscle biopsies), as well as by their limited expansion capacity and premature differentiation *in vitro*, which pose major hurdles to reach the cell number required to treat patients with disorders involving multiple large muscles such as DMD (Cossu *et al*., 2016). Induced pluripotent stem cells (iPSCs) offer a solution to bypass these limitations.

Human iPSCs (hiPSCs) are increasingly becoming a key source of skeletal myogenic progenitor cells for disease modelling and transplantation studies, owing to their controllable proliferation and differentiation capacity, lack of significant ethical concerns and non-invasive sampling of the starting primary cell population (Loperfido *et al*., 2015). Several protocols are currently available to generate skeletal myogenic derivatives from hiPSCs (reviewed in (Selvaraj, Kyba and Perlingeiro, 2019)). Starting from the pioneering studies based upon controlled expression of myogenic regulators to obtain transplantable skeletal myogenic cells from hiPSCs (e.g., Darabi *et al*., 2012; Goudenege *et al*., 2012; Tedesco *et al*., 2012), the field has refined transgene-based protocols to direct hiPSC differentiation into skeletal muscle (e.g., Albini *et al*., 2013; Maffioletti *et al*., 2015; Shoji *et al*., 2016; Kim *et al*., 2021), whilst also developing genomic-integration-free, small molecule-based methods to derive myogenic cells mimicking embryonic development (e.g., Borchin *et al*., 2013; Caron *et al*., 2016; Chal *et al*., 2016; Hicks *et al*., 2018). However, the focus on perfecting methods to obtain myogenic progenitors resembling self-renewing MuSCs has neglected the critical need to enhance their migration capacity, which is essential to deliver cells to large or multiple muscle districts. As a result, no defined methods are currently available to differentiate hiPSCs into myogenic progenitors with high migratory and/or trans-endothelial migration capacity.

Here we describe a developmentally-inspired, small-molecule-based, transgene- and genomic-integration-free strategy to increase motility and trans-endothelial migration of hiPSC-derived myogenic progenitors (hiMPs) via modulation of NOTCH and PDGF signalling, following their specification from pluripotent cells. This study (summarised in figure 1) provides a clinically relevant framework to improve migration of hiMPs for future cell therapies of muscle diseases.

**Figure 1.**
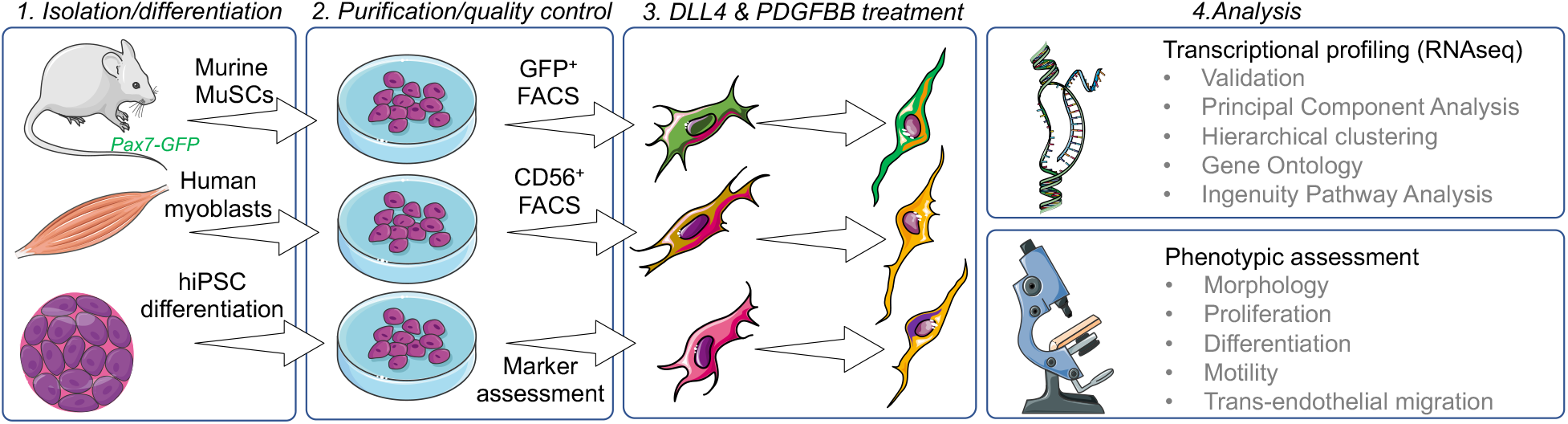
Schematic representation of the cell isolation, culture, treatment, differentiation and analysis pipeline underpinning this study. Figure created using Servier Medical Art (https://smart.servier.com) in accordance with a Creative Commons Attribution 3.0 Unported License (https://creativecommons.org/licenses/by/3.0/).

## RESULTS

### Combined activation of NOTCH and PDGF signalling pathways induce conserved transcriptional changes in mouse and human tissue-and iPSC-derived myogenic progenitors

We aimed to identify targetable pathways to improve migration of hiMPs, and focused on NOTCH and PDGF (Hellström *et al*., 1999; Armulik, Genové and Betsholtz, 2011) which have been shown to improve migration in tissue-resident MuSCs (Gerli *et al*., 2019). To identify whether hiMPs respond to activation of the aforementioned pathways, we performed an unbiased assessment of global transcriptomic changes induced by DLL4 and PDGF-BB in wild-type mouse and human primary MuSC-derived myoblasts, alongside hiMPs. Before performing bulk RNA-sequencing (RNAseq) of these cell populations, we first assessed their purity. Four distinct mouse and four distinct human MuSC-derived myoblast populations were isolated and FACS-purified from skeletal muscles of Pax7-nGFP mice and from normal human muscle biopsies using green fluorescent protein (GFP) and CD56 (NCAM), respectively (Materials and Methods). hiMPs were derived from four different, fully-characterised hiPSC lines generated with genomic-integration-free technologies using a validated transgene-free, small molecule-based protocol recapitulating skeletal muscle developmental specification and differentiation *in vitro* (Caron *et al*., 2016) (Materials and Methods). Purity of the hiPSC derivatives was assessed by immunostaining for myogenic and other non-myogenic markers. All assessed hiMPs were homogeneously positive for the skeletal myogenic determination factor MYOD and lacked contamination from neuroectodermal derivatives (PAX6 and MAP2) (figure S1). After assessing their purity, the three groups of cells were treated for 7 days with DLL4 and PDGF-BB (Materials and Methods) and then RNA was extracted from treated and untreated samples for RNAseq. Principal component analysis (PCA) revealed distinct segregation between DLL4 and PDGF-BB-treated and untreated populations of the 3 cell types (figure 2A, table S1,2). Additionally, RNAseq analysis provided a total of 1405, 337 and 2990 differentially expressed genes between treated mouse MuSC-derived myoblasts, human myoblasts and hiMPs and their untreated controls respectively (figure 2B). Hierarchical clustering of top 50 differentially regulated genes in mouse and human samples showed overall consistency of transcriptional dynamics in those transcripts across all four lines analysed (figure S2, table S2).

**Figure 2.**
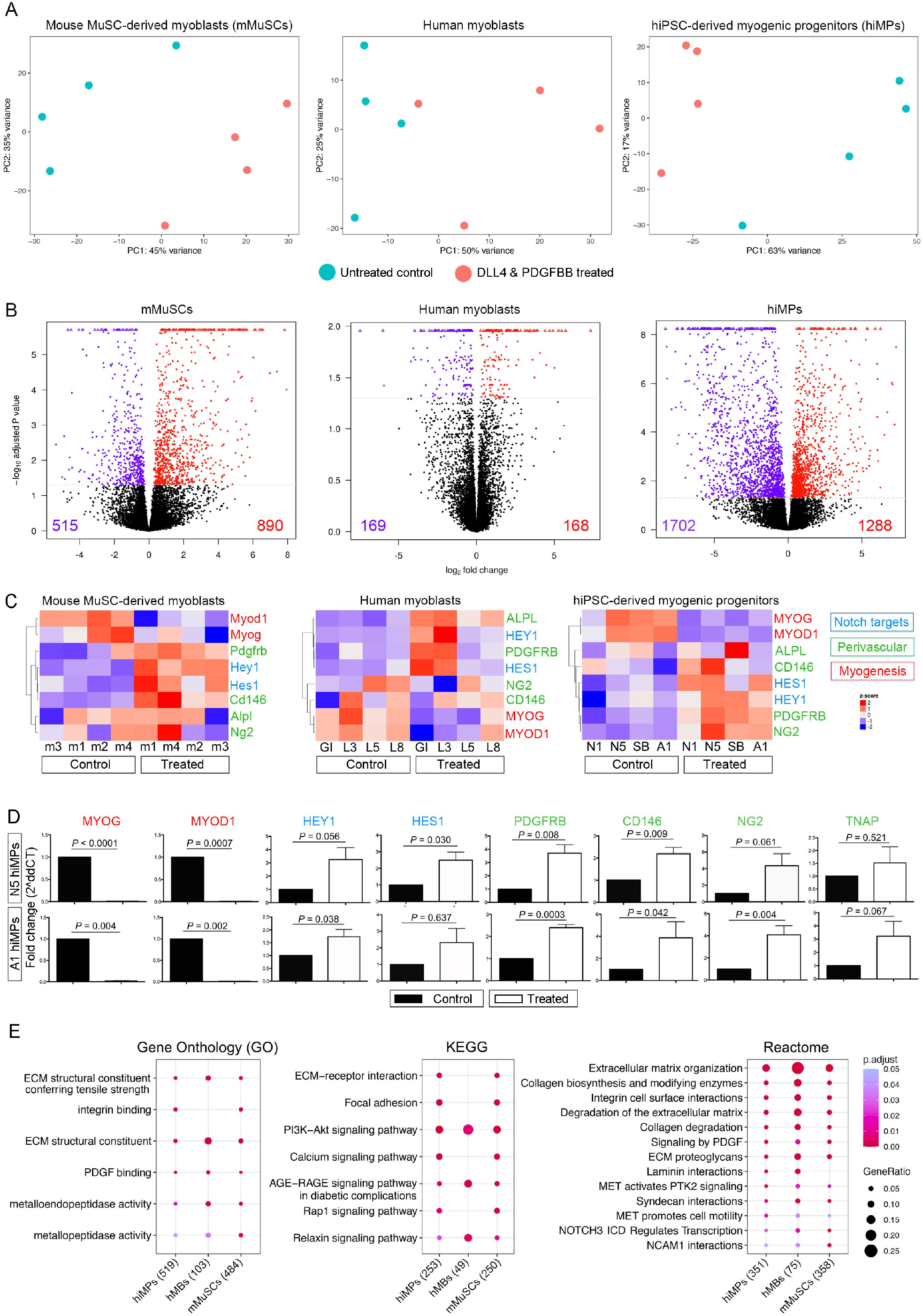
RNAseq-based transcriptional profiling of mouse and human myogenic progenitors upon activation of NOTCH and PDGF signalling pathways. **(A)** Principal Component Analysis (PCA) showing mMuSC-derived myoblasts (left), human myoblasts (centre) and hiMPs (right). 4 cell lines were analysed with RNAseq in treated and untreated conditions for each cell population. Each point on the PCA represents a cell population. Additional information in Table S1,2. **(B)** Volcano plots visualising differentially expressed genes between untreated and DLL4 & PDGFBB-treated mMuSCs, human myoblasts and hiMPs. Red dots represent genes which display a positive fold-change in expression upon treatment with DLL4 & PDGF-BB whilst violet dots represent genes which are significantly downregulated. Differentially expressed genes required a *P* value of ≤ 0.05. **(C)** Heatmaps showing changes in expression of key myogenic (*MYOD, MYOGENIN*), perivascular (*PDGFRB, NG2, CD146, ALPL*) and NOTCH target (*HEY1, HES1*) genes upon treatment with DLL4 & PDGF-BB in mMuSC-derived myoblasts (left), human myoblasts (middle) and hiMPs (right). Clustering was performed by genes/probes with Pearson correlation. Colour scale based on z-scores: red regions indicate high expression whilst blue regions indicate low expression. Dendrograms indicate the similarity of clusters as well as the orders in which clusters were assembled. **(D)** Validation of RNAseq data of panel **(C)** by real-time PCR analysis of the same myogenic, perivascular and NOTCH target transcripts in treated and untreated hiMPs (N=3; error bars; S.E.M.). Statistical analysis (paired *t* test) performed on ΔCt values whilst graphs were produced as fold change relative to untreated controls. **(E)** Curated dot plot Gene Ontology (GO; left), Kyoto Encyclopaedia of Genes and Genomes (KEGG; centre) and Reactome (right) enrichment analyses showing shared gene functions amongst the cell groups; numbers in brackets: genes analysed with a p value threshold set at 0.05; full lists in a dedicated spreadsheet available in Supplemental Information.

We then tested whether the observed global transcriptional changes were a consequence of NOTCH and PDGF signalling activation. To address this question, we looked at specific downstream targets of NOTCH and PDGF pathways, as well as key myogenic and perivascular markers known to be modulated by this treatment in murine myoblasts (Cappellari *et al*., 2013; Gerli *et al*., 2019). As shown in Figure 2C, treated mouse MuSCs (mMuSCs), human myoblasts and hiMPs shared similar dynamics of NOTCH targets and perivascular transcripts upregulation, coupled with downregulation of myogenic transcripts such as *Myogenin* and *MyoD* (also a downstream NOTCH signalling target (Kopan, Nye and Weintraub, 1994)). This was further validated via qRT-PCR analysis in two representative hiMP lines (figure 2D). We subsequently wanted to identify inter-species similarities in transcriptional response to DLL4 and PDGF-BB treatment. For this purpose, we selected the top 50 differentially regulated genes of treated mMuSC-derived myoblasts, found the relative human orthologues and then performed supervised hierarchical clustering on human myoblast and hiMP datasets. The resulting heatmaps show that the majority of transcripts in the treated human cells display a similar regulation in comparison to their murine counterparts (figure S3; table S3), further indicating an overall conservation of the cellular response to DLL4 and PDGF-BB in skeletal myogenic cells (albeit with some expected variability in human, non-syngeneic cells). Finally, Gene Ontology (GO), Kegg and Reactome enrichment analyses showed shared gene functions amongst the cell groups, including pathways involved in extracellular matrix remodelling, integrin-cell surface interactions, focal adhesion generation, in addition to the expected NOTCH and PDGF pathways (figure 2E).Together these data demonstrate that DLL4 and PDGF-BB induce transcriptional changes across skeletal myogenic progenitors from different species and developmental origins, with hiMPs showing the greatest transcriptional response.

### Analysis of morphology, proliferation and differentiation of DLL4 and PDGFBB-treated hiMPS

To identify if the transcriptional response of DLL4 and PDGFBB-treated hiMPs results in detectable, cellular phenotypic changes, we assessed specific transcriptional signatures alongside functional readouts such as morphology, proliferation and skeletal myogenic differentiation capacity. Hierarchical clustering analysis highlighted modulation of several regulators of cell morphology such as upregulation of *MYH9, MYO10, RAC1/3* and *RHOC*, alongside downregulation of *RHOD, MYH10, ITGA7* and *SEMA3* (figure 3A; table S4). After one week of treatment, hiMPs appeared more elongated than their untreated counterpart, in accordance with what was observed in mMuSCs (Gerli *et al*., 2019). Morphometric analysis confirmed this finding, revealing a higher number of cells falling within the first quartile (0-0.25) of cell circularity ratio (i.e. cells with marked protrusions; figure 3B-C; mean ± SD: treated 45.33 ± 10.26, untreated 12.67 ± 10.60; *P* = 0.027, paired *t*-test). We next assessed the impact of DLL4 and PDGF-BB treatment on hiMP proliferation. A decrease in the proliferative capacity of myogenic cells could be detrimental for cell therapy, limiting the translational potential of donor cells. Hierarchical clustering analysis highlighted modulation of several regulators of cell proliferation and lineage commitment in at least 3 out of 4 hiMP lines, including upregulation of *PDGRB, NOTCH3, VEGFA* and *TGFB1*, alongside downregulation of *CTNNBIP1, HMGB2* and the myogenic factors *MEF2C* and *MYOG* (figure 3D). These transcriptional changes did not impact on the proliferative ability of hiMPs, with treated and untreated cells displaying a comparable cell cycle, as shown by functional EdU incorporation assay (figure 3E-F; table S4).

**Figure 3.**
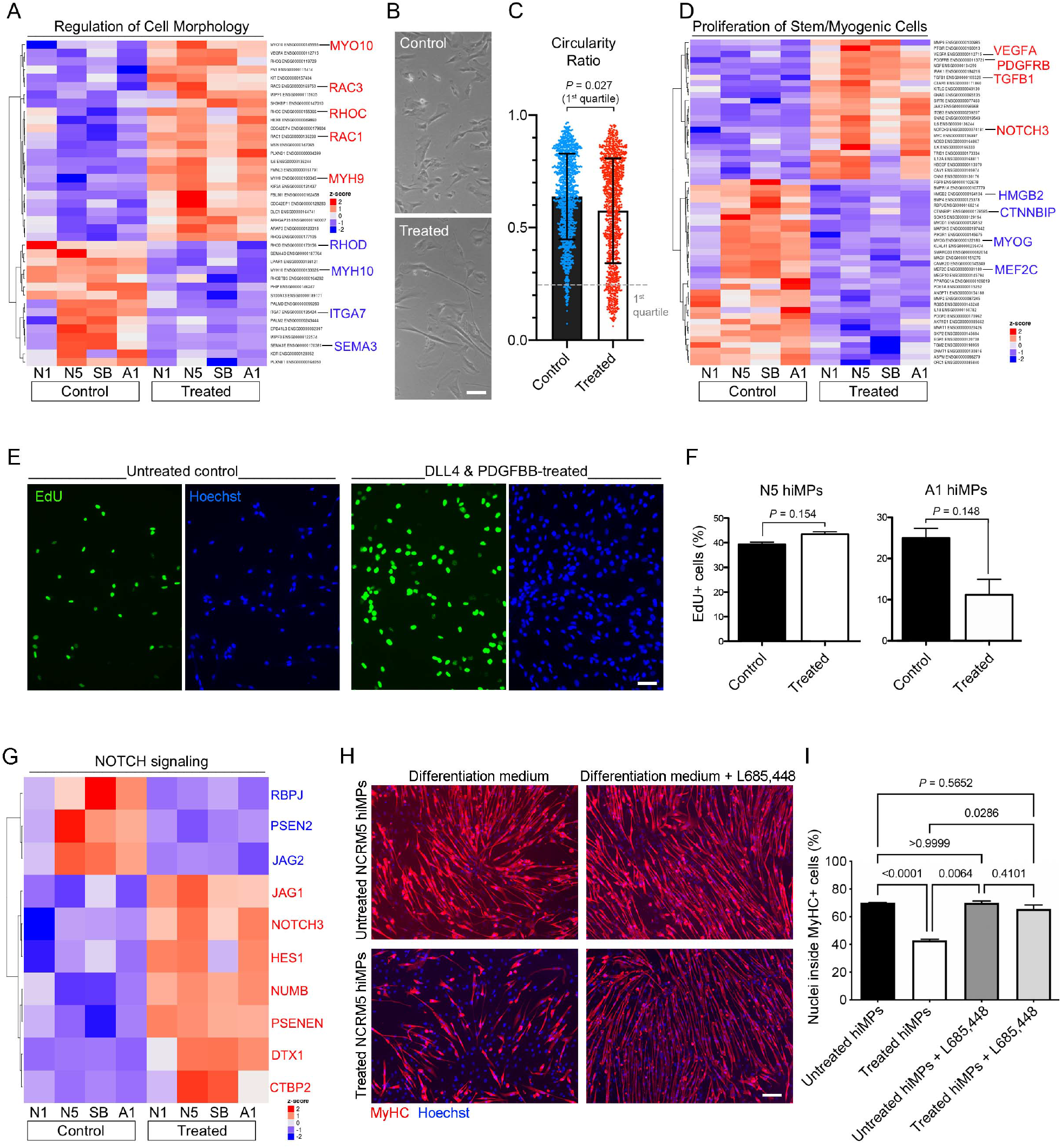
Analysis of morphology, proliferation and differentiation of DLL4 and PDGFBB-treated hiMPs. **(A)** *P* value-adjusted hierarchical clustering heatmap generated from a gene ontology list of genes involved in regulation of cell morphology (GO 0008360; *P* set at 0.05). **(B)** Phase contrast images displaying morphology of untreated and treated hiMPs. Scale bar: 25 μm. **(C)** Scatter dot plot showing morphometric analysis of treated and untreated hiMPs. Morphology was quantified as circularity ratio, where 1 = perfect circle and 0 = line (N = 3). Statistical analysis (paired *t* test) was performed on the first quartile to enhance detection of morphological changes (error bars: SD). **(D)** *P* value-adjusted hierarchical clustering heatmap generated from a gene ontology list of genes involved in proliferation of stem and myogenic cell types (GO 2000291; 0048660; 0014857; 0072091; *P* set at 0.05). **(E)** Immunofluorescence images of untreated and treated hiMPs incubated with EdU for 2 hours. Scale bar: 75 μm. **(F)** Bar graphs quantifying EdU experiment shown in (E) (N = 3; error bars: SEM). **(G)** *P* value-adjusted hierarchical clustering heatmap of NOTCH signalling genes (Kegg pathway 04330). **(H)** Immunofluorescence images of hiMPs expanded in control or treated conditions for 1 week, induced to differentiate for 4 days in the presence or absence of γ-secretase inhibitor L685,448 and immunostained for myosin heavy chain (MyHC). **(I)** Bar graph quantifying the average percentage of nuclei within MyHC positive myotubes (N = 3; error bars: SEM). Statistical significance based on one-way ANOVA with Tukey’s multiple comparison. Scale bar: 75 μm. Full gene list for heatmaps in (A) and (D) available in Table S4.

NOTCH activation inhibits myogenesis *in vitro* in embryonic and adult myoblasts (Kopan, Nye and Weintraub, 1994; Conboy and Rando, 2002; Mourikis and Tajbakhsh, 2014; Gerli *et al*., 2019). Although RNAseq analysis of the NOTCH pathway shows modulation of several effectors (figure 3G), we wanted to verify functionally if this phenomenon was also conserved in hiMPs. To achieve this aim, we induced myogenic differentiation of DLL4 & PDGF-BB treated cells and observed a significant reduction in the percentage of nuclei within MyHC-positive fibres, from 70.00% ± 0.29% to 43.06% ± 1.20% (figure 3H-I; *P*<0.0001; N = 3; mean ± SD; one-way ANOVA with Bonferroni correction). To further validate the NOTCH-dependency of this finding, we blocked NOTCH pathway with the γ-secretase inhibitor L685458, which selectively inhibits γ-secretase-dependent nuclear translocation of the NOTCH Intra-Cellular Domain (NICD) (Vilquin *et al*., 1994). Upon treatment with L685,448, the impairment of differentiation was reverted from 43.06 ± 1.20 to 65.59 ± 5.11 (*P* 0.028; figure 3H-I), thus confirming that hiMP myogenic differentiation potential is NOTCH-dependent and could be restored to pre-treatment levels.

### Combined DLL4 and PDGF-BB treatment enhances migration of hiMPs

We and others have shown that NOTCH and PDGF pathways play a critical role in regulating developmental fate, regenerative potential and migration of primary, native myogenic cells (Betsholtz, 2004; Cappellari *et al*., 2013; Piñol-Jurado *et al*., 2017; Camps *et al*., 2019; Gerli *et al*., 2019). Of the aforementioned properties, cell migration is of key relevance for cell therapy. To investigate if DLL4 and PDGF-BB had an effect on cell migration of hiMPs, we analysed the differentially expressed genes in our RNAseq dataset using Ingenuity Pathway Analysis (IPA). Amongst the most significantly modulated cellular functions upon DLL4 & PDGF-BB treatment there was “Cellular Movement”, with a total of 578 differentially expressed genes (figure 4A). To correlate these transcriptional changes to a phenotypic response, single cell tracking of cells exposed to DLL4 & PDGF-BB was performed using a deep learning approach in an automated manner and track features were extracted using Heteromotility (Kimmel *et al*., 2018) (figure 4B-E; Materials and Methods). Visualisation with t-SNE plots revealed that both untreated and DLL4 & PDGF-BB-treated hiMPs shared a similar motility profile (figure S4A). To identify heterogenous motility phenotypes, unsupervised hierarchical clustering (Ward’s method) was performed, which captured >95% variation within the system to obtain two clusters (Silhouette *Si* = 0.19) (Figure 4C and S4B). Cluster 1 comprises a less motile population of cells as indicated by lower total distance travelled, average speed and average time spent moving (Figure 4D). Cells within cluster 1 also display higher non-Gaussian parameter and kurtosis which suggests levy flight-like motions, a classical model of motion (figure 4D). Cluster 2 represents the motile population of cells, demonstrating higher distances travelled, average speeds and proportion of time spent moving. Additionally, cells within cluster 2 performed directed migration as shown by higher progressivity, linearity and mean squared displacement (MSD) (figure 4D). Notably, a higher proportion of treated cells was observed within cluster 2 in comparison to untreated controls (Figure 4E, S4). Furthermore, analyses performed with Trackmate (Materials and Methods) validated these findings, showing increased trends in distance, straight line speed, progressivity and velocity in treated hiMPs (figure S4E). To have further insights on possible protein-protein interaction networks that might positively regulate cell migration, we subsequently analysed our RNAseq dataset with the STRING platform (https://string-db.org; (Szklarczyk *et al*., 2019); figure 4F). Functional enrichment analysis highlighted that a number of candidate proteins with relevance in cell migration, which could be associated with the observed migratory phenotype, were upregulated in our datasets, such as TGFB1, ADAMTS2/12/14 and THY1 (Figure 4F) (Barker *et al*., 2004; Sciorati *et al*., 2006; Li *et al*., 2020).

**Figure 4.**
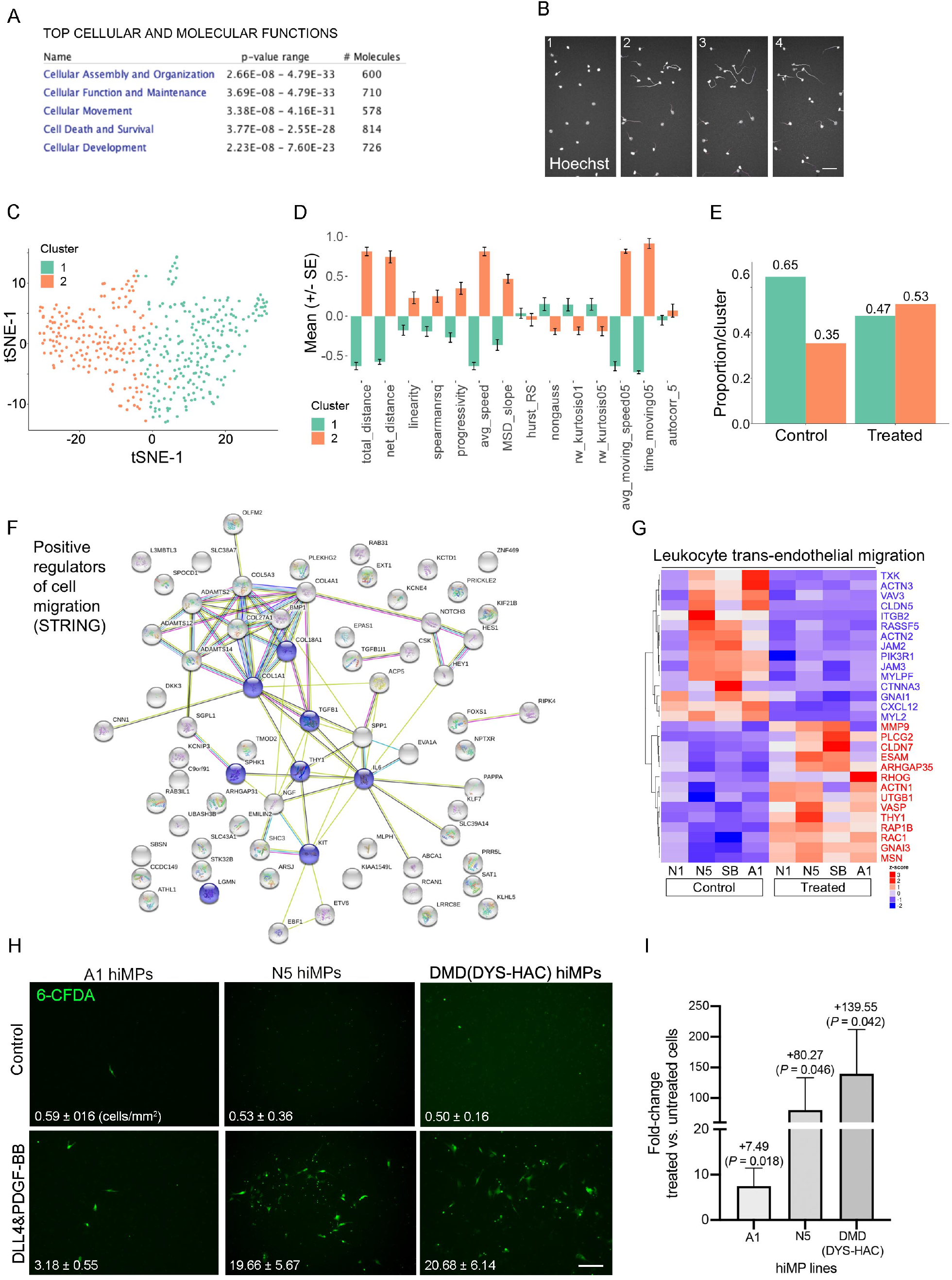
Transcriptional and functional analyses of the DLL4 and PDGFBB treatment on motility and migration of hiMPs. **(A)** Top cellular and molecular functions associated with DLL4 & PDGFBB modulation generated via ingenuity pathway analysis (IPA). Genes upregulated in the DLL4 & PDGFBB-treated hiMPs relative to the untreated control were subjected to IPA to reveal the predicted most significant associated functions. **(B)** Fluorescence microscopy images depicting Hoechst-positive nuclei of each cell at sequential time points. Coloured tails represent the locations of the nuclei at previous time points. **(C)** Unsupervised hierarchical clustering (Ward’s method) visualised with a t-SNE plot showing two distinct clusters (Silhouette *Si* = 0.19) (n = 408) (perplexity = 35). Cells pooled from 3 independent experimental replicates for each condition were used (untreated and DLL4 & PDGF-BB-treated). **(D)** Bar chart demonstrating normalised values for comparison of motility phenotypes between cells within the two clusters (mean ±SEM). Statistical significance based on Bonferroni-corrected *t*-test: all parameters except “hurst_RS” and “autocorr” are statistically significant between the two groups p<0.01 (data points: single cells pooled together from 3 independent experiments). **(E)** Bar graph displaying the proportions of control and DLL4 & PDGF-BB-treated cells within each cluster. **(F)** Functional protein association network analysis (https://string-db.org). The network view summarises predicted associations for proteins positively regulating cell migration common to all three datasets. The nodes are proteins and the edges represent the predicted functional associations. Red line: fusion evidence; Green line: neighbourhood evidence; Blue line: co-occurrence evidence; Purple line: experimental evidence; Yellow line: text mining evidence; Light blue line: database evidence; Black line: co-expression evidence. Blue nodes: GO:0030335 positive regulation of cell migration, Count in gene set: 8 of 452, false discovery rate: 0.0156. **(G)** *P* value-adjusted hierarchical clustering heatmap displaying hierarchical clustering of genes associated with leukocyte trans-endothelial migration (KEGG pathway: hsa04670; *P* set at 0.05). **(H)** Assessment of DLL4 & PDGF-BB-treated WT and genetically corrected DMD hiMP migration through a layer of endothelial cells. Representative images showing the lower side of the trans-well membrane on which treated and untreated hiMPs (stained with the transient dye CFDA, in green) are simultaneously seeded on HUVECs for 8 hours. Bar graphs quantifying the average number of CFDA-positive cells/ mm^2^, that have migrated through the endothelial layer in each considered condition. (N = 3). A minimum of 10 (1.5 mm^2^) fields per condition was quantified (mean ±SEM). Scale bar: 250 μm. **(I)** Bar graph showing fold-change in trans-endothelial migration (mean ±SEM). Statistical significance based on one-way ANOVA with Bonferroni’s multiple comparison.

Although encouraging, enhanced cellular motility may not be directly relevant in the context of cell therapies requiring intravascular cell delivery to target multiple large muscles. Therefore, we assessed the effect of DLL4 and PDGF-BB treatment on trans-endothelial migration which, conversely, is essential for systemically delivered cells. Interestingly, RNAseq analysis showed positive modulation of several transcripts involved in cell extravasation in treated hiMPs, such as *ESAM, ICAM3, JAM2, MMP9, PDGFD* and *THY1*, although some other mediators of extravasation such as *ITGB2* and *CXCL12* were downregulated (Figure 4G, S4; Table S4, S6). To functionally assess the trans-endothelial capacity of DLL4 and PDGF-BB treated hiMPs we performed trans-well migration assays. After 7 days of treatment with DLL4 and PDGF-BB, hiMPs were incubated with a transient fluorescent dye (CFDA, Materials and Methods) and plated onto a monolayer of human-umbilical vein endothelial cells (HUVECs). After 8 hours membranes were fixed and CFDA-positive, trans-migrated cells were quantified. As shown in Figure 4H-I, treatment with DLL4 and PDGF-BB significantly enhanced the ability of hiMPs to migrate through an endothelial monolayer (from 0.59 cells/mm^2^ to 3.19 cells/mm^2^ (*P* = 0.0180) and from 0.50 cells/mm^2^ to 20.68 cells/mm^2^ (*P* = 0.0464), in healthy donor hiMPs, respectively). Similar results were obtained with hiMPs derived from a DMD patient and genetically-corrected with a human artificial chromosome containing the entire 2.5Mb *DYSTROPHIN* genetic locus (DYS-HAC, Materials and Methods)(Choi *et al*., 2016), demonstrating that even after genetic correction, hiMPs remain responsive to the DLL4 and PDGF-BB treatment (Figure 4H-I). Collectively, these results show that modulation of NOTCH and PDGF signalling pathways improves migration of hiMPs across endothelial monolayers, laying the foundation for future studies aimed at elucidating and further enhancing the molecular mechanism underpinning this phenomenon.

## DISCUSSION

Here we report a transgene-free strategy to confer transmigration properties to hiMPs, showing that DLL4 and PDGF-BB treatment of hiMPs induces a transcriptional profile comparable to those detected in MuSCs from mouse and human primary samples, including key markers of myogenic commitment, downstream NOTCH signalling targets and perivascular markers. This transcriptional response is most likely caused by the role of DLL4 and PDGF-BB as developmental determinants of skeletal muscle pericytes (Cappellari *et al*., 2013; Moyle, Tedesco and Benedetti, 2019). Notably, hiMPs responded to the treatment more consistently than adult mouse and human myoblasts, possibly due to their relatively immature state compared to their adult counterpart. Interestingly a recent study showed that hiMPs are transcriptionally comparable to late embryonic and early foetal myoblasts (Xi *et al*., 2020). Secondary myogenesis occurs between E14.5-17.5 of mouse development during which foetal myoblasts either contribute to existing primary myofibres or fuse with each other to give rise to secondary muscle fibres (Messina and Cossu, 2009). Spontaneous fate transitions of myoblasts to pericytes were observed in foetuses at E16.5 (Cappellari *et al*., 2013). Therefore, it is possible that hiMPs reflect this plastic foetal myoblast nature and therefore might respond more robustly to DLL4 and PDGF-BB treatment.

Morphological analysis between treated and untreated hiMPs revealed that DLL4 and PDGF-BB induced shape changes in a subset of treated cells, in keeping with perturbations of the actin cytoskeleton highlighted by RNAseq data (e.g., genes of pathways differentially expressed include *Actin cytoskeleton, Signalling by Rho GTPases*, and *Actin nucleation by ARP-WASP complex*, data not shown). Several studies have demonstrated that chemokines enhance myoblast migration via direct regulation of the actin cytoskeleton (Kawamura *et al*., 2004; Ishido and Kasuga, 2011). For example, hepatocyte growth factor-mediated increase of migration is facilitated by lamellipodia formation via the PI3K/AKT and ERK/MEK signalling pathways (González *et al*., 2017). Additionally, stromal-derived factor 1 (SDF1), via interaction with CXCR4, increases migration via upregulation of Rho GTPases, *CDC42* and *Rac1* in addition to several other migration-associated transcripts such as actin bundling protein, *ACTN1* and calcium-dependent protease, *CAPSN1*, necessary for cleavage of focal adhesions (Kowalski *et al*., 2017). The aforementioned presence of a subset of cells displaying pronounced morphological changes in response to treatment could also indicate the existence of multiple cell states within the hiMP population with differential susceptibility to perturbations of NOTCH and PDGF signalling, also suggested by our migration analyses. Future work should consider correlating single cell RNAseq and motility analyses to identify responders, characterising the aforementioned cell states, increasing purity of the treated population and defining molecular mechanisms responsible for increased cell migration.

Any viable treatment to improve engraftment of cell therapy products should not impact negatively on their proliferation and differentiation capacity. DLL4 and PDGF-BB treatment did not alter hiMP proliferation and the expected NOTCH-mediated reduction in differentiation was fully rescued by γ-secretase inhibition of NOTCH signalling. Although future studies may be needed to elucidate the requirement for γ-secretase treatment pre-transplantation, it is likely that a spontaneous re-acquisition of skeletal myogenic capacity will occur *in vivo* upon removal of the exogenous DLL4 stimulus, as previously reported (Gerli et al., 2019).

*In silico* analyses and *in vitro* assays indicated that treated hiMPs possess enhanced motility and trans-endothelial migration, validating our initial hypothesis that DLL4 and PDGF-BB modulate migration also in hiPSC derivatives. These findings prompted us to explore relevant signalling pathways and molecules involved in this process (reviewed in Choi, Ferrari and Tedesco, 2020). Leukocytes are the typical benchmark for extravasating cells and many genes regulating leukocyte extravasation were found to be differentially expressed upon DLL4 and PDGF-BB treatment in hiMPs. Interestingly, certain molecules relevant for leukocyte extravasation such as CXCL12 and integrin β2, were downregulated upon DLL4 and PDGF-BB treatment, suggesting that myogenic progenitor trans-endothelial migration may not necessarily mimic all aspects of leukocyte extravasation. For example, downregulation of *JAM-A* enhances trans-endothelial migration of adult muscle pericyte-derived mesoangioblasts (Bonfanti *et al*., 2015). Furthermore, intra-arterial delivery of adult mouse mesoangioblasts in *JAM-A-null* dystrophic mice resulted in increased engraftment, indicating that absence of endothelial JAM-A improves trans-endothelial migration of myogenic cells (Giannotta *et al*., 2014). This is in contrast to leukocytes, in which isophilic interactions between JAM-A of leukocytes and endothelial cells are necessary for efficient extravasation (Corada *et al*., 2005; Woodfin *et al*., 2007, 2009). Therefore, although not all mechanisms of leukocyte extravasation are mimicked by myogenic progenitors, it remains possible that conserved elements may exist (e.g. figure 4G). Combining these features with the machinery that other non-haematopoietic cells use to travel through endothelia (e.g., metastatic cancer cells) could provide additional tools for myogenic cells to efficiently extravasate. Future studies should also investigate *in vitro* high-throughput and high-fidelity methods to evaluate cell transmigration, ideally using organotypic (i.e., skeletal muscle-specific) endothelial and smooth muscle cells on top of a basement membrane, which could facilitate unravelling of adhesion profiles and tissue-specific recruitment mechanisms necessary for efficient trans-endothelial migration. This strategy may be more informative than interspecific *in vivo* experiments based upon hiMPs delivery within murine blood vessels, where the species mismatch could affect receptor recognition and downstream signalling (e.g., limited interactions between human-mouse selectins/integrins). In summary, this study provides an important first step towards the identification of druggable targets to increase the migration capacity of hiMPs, ultimately contributing to the identification of a systemically deliverable and engraftable hiPSC-derivative for muscle cell therapies.

## MATERIALS AND METHODS

### Cell isolation and culture

Primary mMuSCs were isolated, purified via FACS from skeletal muscles of four distinct *Tg:Pax7-nGFP* F1:C57BL/6:DBA2 mice expressing nuclear-localized EGFP in Pax7-expressing cells (Sambasivan *et al*., 2009) and cultured as previously reported (Gerli *et al*., 2019). C2C12 myoblasts (Yaffe and Saxel, 1977) were cultured in DMEM supplemented with 10% FBS supplemented with 10% FBS, 1% glutamine (Sigma-Aldrich) and 1% penicillin-streptomycin. Primary human myoblasts from three different donors were obtained from the MRC Neuromuscular Centre Biobank (shortened as L3, L5 and L8), FACS-purified for CD56^+^ (Biolegend; CD56-FITC 304604) and cultured as previously reported (Gerli *et al*., 2019); an additional polyclonal population of biopsy-derived human myoblasts was purchased (Gibco human skeletal myoblasts A12555, shortened in the text as GI) and purified using CD56. Five different hiPSC lines have been used in this study; four lines have been used for most experiments: N1 (short for NCRM-1: https://hpscreg.eu/cell-line/CRMi003-A), N5 (short for NCRM-5: https://hpscreg.eu/cell-line/CRMi001-A); SBI (short for SBIi006-A: https://hpscreg.eu/cell-line/SBIi006-A); A1 (short for: Gibco Episomal hiPSC line A13777); Genetically-corrected DMD(DYS-HAC) iPSCs were generated using Sendai-virus-delivered reprogramming factors and kindly provided by Dr. Y. Kazuki and Prof. M. Oshimura (Tottori University, Japan; (Choi *et al*., 2016)). hiPSCs were cultured on vitronectin XF^™^ (Stemcell technologies; 07180) at 37°C, 5% CO2 and 3% O2. mTESR-E8^™^ (Stemcell technologies; 07174) was used for cell expansion and colonies were passaged using gentle dissociation media (STEMCELL technologies; 07174) according to manufacturer’s instructions. Skeletal myogenic differentiation of hiPSCs was performed with a commercially available protocol (Myocea) (Caron *et al*., 2016). Briefly, hiPSCs were dissociated into single cells and plated onto Matrigel-coated (Corning) dishes. Subsequently, cells were exposed to induction medium for 10 days to generate myogenic progenitors. The resulting hiMPs were expanded in myoblast growth and proliferation media (Lonza; SKBM-2). All myogenic cells were differentiated in DMEM containing 2% horse serum. Human cell work was conducted under the approval of the NHS Health Research Authority Research Ethics Committee reference no. 13/LO/1826; IRAS project ID no. 141100.

### DLL4, PDGF-BB and γ-secretase inhibitor treatment

Recombinant human DLL4 (DLL4 fused with the Fc domain of human IgG; R&D Systems; 1506-D4) was resuspended to a final concentration of 10 μg/ml in sterile PBS containing 1% wt/vol bovine serum albumin (BSA; Sigma-Aldrich; A9418-10G) as a carrier protein. Standard cell culture plastic dishes were coated with the DLL4 solution and incubated at 37° C for 45 minutes. Cells were then seeded on the coated flasks and supplemented with 100 ng/ml of human PDGF-BB re-suspended in 0.1%BSA/4mM HCl/PBS (R&D Systems; 200-BB-050) daily for at least 7 days. As for normal myogenic differentiation assays, DLL4 and PDGF-BB-treated myoblasts and untreated control were seeded at a high density on collagen-coated dishes and, when confluent, switched to differentiation medium (DMEM supplemented with 2% HS (w/v) + 1% P/S (w/v)). To block NOTCH signalling cells were incubated with 660 ng/ml of *γ*-secretase inhibitor (L685, 448, Sigma) 24 hours before the switch to differentiation medium and over the two following days.

### RNA sequencing

#### RNA library preparation

For RNA-seq analyses, hiMPs were differentiated from the four aforementioned hiPSC lines using the previously described Myocea kit (Caron *et al*., 2016). Four primary human biopsy-derived myoblast populations were obtained from the UCL MRC Neuromuscular Centre Biobank (L397, L588 and L876) as well as from pooled populations of biopsy-derived cells (Gibco) and subsequently FACS-purified for CD56. Primary mouse myoblasts were isolated via FACS for GFP from skeletal muscles of *Pax7-nGFP* mice as previously reported (Sambasivan *et al*., 2009). Cells were seeded on dishes coated with 10ug/ml DLL4 and medium supplemented daily with 50ng/ml recombinant PDGF-BB with a minimum of 1 passage throughout the 7-day reprogramming protocol to replace the DLL4 protein. An untreated control was grown in parallel on 1% BSA-coated dishes. After 7 days, samples were collected and RNA extracted with Qiagen RNeasy kit, with on-column DNaseI treatment. RNA concentration and integrity were assessed by Nanodrop spectrophotometer and Agilent 2100 bioanalyzer (model G2939A). An RNA Integrity Number (RIN) (Schroeder *et al*., 2006), was quantified for each sample and scores between 9.8-10 accepted. Library preparations were performed with the UCL Genomics facility, using the KAPA mRNA HyperPrep Kit (Roche) to capture mRNA and deplete ribosomal RNA. Samples were barcoded and run together on an Illumina NextSeq 550 System to minimise batch variation.

#### Analyses

Raw sequence data were pre-processed to remove small (>20bp) or poor quality reads using Trimmomatic v0.36.4 (Bolger, Lohse and Usadel, 2014). Reads were aligned either to the Human hg38 genome using Spliced Transcripts Alignment to a Reference (STAR) software v2.5.2b (Dobin *et al*., 2013), mapped reads de-duplicated with Picard v2.7.1.1 (Broad Institute) and reads-per-transcript calculated with feature Counts v1.4.6.p5 read summarization tool (Liao, Smyth and Shi, 2014). Finally, differential expression was calculated using SARTools R package v.1.3.2.0 (Varet *et al*., 2016), based on the DESeq2 model and package (Love, Huber and Anders, 2014).

In order to perform gene set enrichment analyses, mouse gene symbols were first converted into their respective human orthologs using the BiomaRt v2.46.2 package (Smedley *et al*., 2009). Subsequently, HUGO Gene Nomenclature Committee (HGNC) symbols were converted into Entrez Gene IDs using BiomaRt. Differentially expressed genes with a fold-change >2 and a *P*-value <0.05 were then subjected to gene set enrichment analysis with ClusterProfiler v3.18.0 and ReactomePA v1.34.0 packages (Yu *et al*., 2012; Yu and He, 2016); script available as supplemental file.

To analyse signalling pathway changes in response to DLL4 & PDGF-BB, differential expression data were inputted into Ingenuity Pathway Analysis (IPA, Qiagen). The Genebank gene ID, log2 fold change expression, p-value and adjusted p-values (padj) were included, in order to account for the experimental false discovery rate. To ensure only highly likely interactions were accounted for, only experimentally observed interactions in mammalian cells were included, and cut-offs were set at log2 fold change (−0.58, +0.58 i.e. a fold change of 1.5) with a padj of 0.05. From this 2259 genes (1002 increased, 1557 decreased) remained on the hiMP dataset. Additional expression analyses were performed using Stemformatics (www.stemformatics.org/) (Choi *et al*., 2019). Functional protein association network analysis was performed using https://string-db.org. RNAseq reads and scripts utilised for PCA and gene enrichment analyses available upon request to the corresponding author. European Nucleotide Archive (ENA) study accession number: PRJEB43338.

### Quantitative real-time PCR

Cells were seeded on 6- or 12-well plates for at least 24 hours before detaching them and centrifuging at 1200rpm to obtain pellets for RNA extraction using RNeasy Mini kit (Quiagen, 74104) according to manufacturer’s instructions. RNA purity and yield were assessed using a Nanodrop spectrophotometer. Retro-transcription to cDNA was performed with the ImProm-IITM Reverse Transcription System kit (Promega, A3800) following manufacturer’s instructions; a minimum of 50 ng of RNA per reaction was used. qRT-PCRs were performed with the SYBR-Green Real Time Master Mix (Promega; A600A) according to manufacturer’s instructions using a BioRad CFX96 machine. qRT-PCRs were performed in triplicate on samples from at least three independent experiments. Ct data were normalised to GAPDH (Stern-Straeter *et al*., 2009). Data were presented as mean ± SEM of the fold change. Significance was assessed on the delta Ct values using student’s two-tailed t-test. List of primers used:

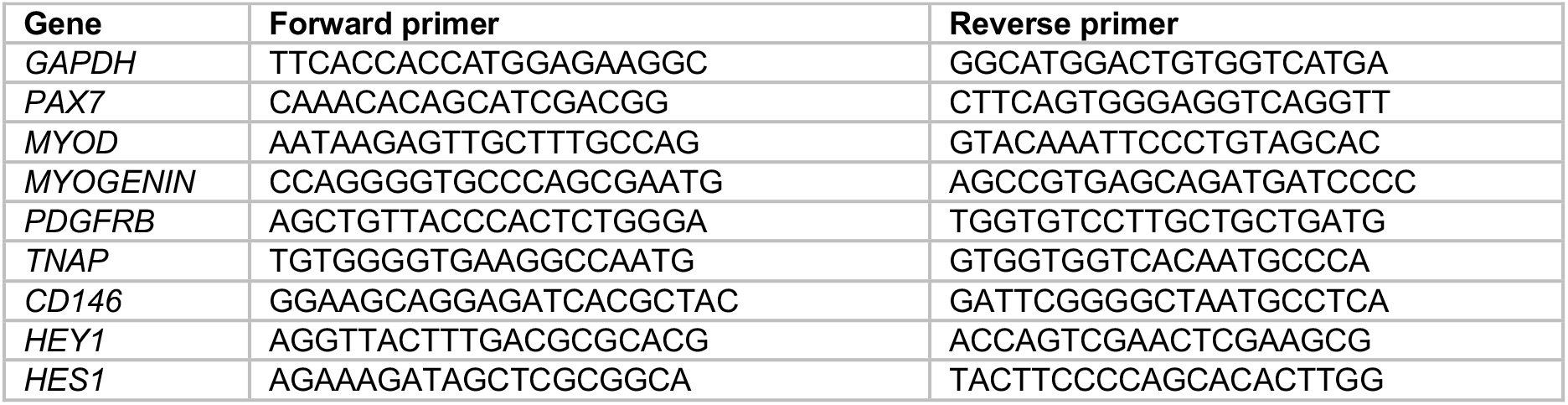

### Immunofluorescence

Cells were fixed in 4% paraformaldehyde (PFA) for 5 minutes at room temperature (RT), washed twice with phosphate buffered saline (PBS) and incubated 30 minutes with PBS-1% BSA-0.2% triton. Cells were then incubated for 30 minutes with 10% donkey or goat serum solution at RT to reduce non-specific antibody binding. Primary antibodies were diluted to the appropriate concentration in PBS-1%BSA-0.2% triton and incubated either one hour at RT or overnight at 4°C. Subsequently, cells were washed three times with PBS-0.2% triton to eliminate unbound antibody and then incubated for 1 hour with fluorescently conjugated secondary antibodies raised in goat or donkey and Hoechst 33342 to visualise nuclei (Fluka; B2261). Cells were imaged using an inverted fluorescence microscope (Leica DMI6000B). At least 5 non-overlapping random field images were acquired and analysed using ImageJ or Adobe Photoshop software.

### Fluorescence Activated Cell Sorting (FACS)

Cells were prepared for FACS analysis as previously published (Maffioletti *et al*., 2015). Briefly, cells were trypsinised and filtered through a 40μm cell strainer in order to get a single cell suspension. At least 1.5 × 10^5^ cells were stained for each fluorochrome-conjugated primary antibody for 1 hour on ice. An additional unstained control tube was included for each cell line. Cells were then washed, fixed in 2 % (w/v) paraformaldehyde for 5 minutes after which 3 mL of FACS buffer was added and cells centrifuged at 232g for 5 minutes. Supernatant was discarded and cells were resuspended in 100 μL FACs buffer and sorted with a CyAn™ ADP Analyser (Beckman Coulter, Inc.) at the UCL GOSICH Flow Cytometry Core Facility. A minimum of 20,000 events per antibody were analysed. FACS data analysis was done using FCS Express 4 (De Novo Software). A similar procedure was followed for FACS cell purification, apart from fixation. Cells were sorted using a MoFlo XDP machine (Beckman Coulter).

### Morphometry and proliferation analyses

To compare morphology between DLL4 and PDGF-BB-treated and untreated hiMPs, the circularity ratio of cells was analysed using ImageJ (0 = line; 1 = perfect circle). Circularity ratios of cells were obtained via quantification of manually labelled cell contours of phase-contrast images. 3 random fields were analysed for 3 independent experiments with at least 300 cells analysed for each biological replicate. To identify differences in proliferation between DLL4 and PDGF-BB-treated and untreated hiMPs, cells were pulsed with 10µM 5-Ethynyl-2’deoxyuridine (EdU) for 2 hours following manufacturer’s instructions (Click-iT® EdU Alexa Fluor® Imaging Kit, Life Technologies). Cells were subsequently fixed and stained with Hoechst 33342. The proportion of proliferating cells was then calculated by comparing the number of EdU+ nuclei with the total number of nuclei within the field.

### Motility and migration assays

1.5 × 10^4^ hiMPs were plated in triplicate onto 24-well multi-well dishes and incubated overnight. Cells were pulsed with Hoechst 33342 (100ng/ml) for 45 mins prior to imaging to aid segmentation and tracking. Imaging was performed with ImageXpress acquiring images every 10 minutes for 12 hours (segmented using ImageJ). Cell tracking, calculation of total distance travelled (μm), velocity (μm/min), mean straight line speed (μm/min) and total displacement (μm) was performed with Trackmate (Tinevez *et al*., 2017). Only cells that remained within the field were analysed. Statistical analysis was performed on three independent repeats with a minimum of 20 cells/condition/repeat.

For analysis of cell motility with Heteromotility, image segmentation was performed using a deep learning approach. A U-net was trained on manually annotated images obtained from individual frames of the tracking dataset selected to capture variation in intensity, shape and size between nuclei. A pre-built U-net was utilised for these purposes (Caicedo *et al*., 2019)(https://github.com/carpenterlab/2019_caicedo_cytometryA) and images were annotated with Caliban (https://github.com/vanvalenlab/caliban). Single-cell tracking was performed with Bayesian Tracker with modified configurations to optimise tracking for videos obtained from ImageXpress (Bove *et al*., 2017) (https://github.com/quantumjot/BayesianTracker). Analysis of single-cell tracks of lengths >60 was subsequently performed with Heteromotility (Kimmel *et al*., 2018) (https://github.com/cellgeometry/heteromotility). The following parameters were utilised: “total_distance” travelled by the cell during time-lapse; “net_distance” travelled by the cell during time-lapse; “linearity”: linear regression analysis of the XY coordinates of a cell at each time point; “spearmanrsq”: assessment of the monotonic relationship of the distribution of XY coordinates of cells at each time point; “progressivity”: ratio between “net_distance” and “total_distance”, serving as an indicator of the directional nature of the cell track during time course. Larger values suggest directional motility; “max_speed”, “min_speed” and “avg_speed”: maximum/minimum/average speed of a cell during time-lapse; “MSD_slope”: spatial deviation of a cell with respect to a reference position during time-lapse. Higher values suggest directional motility whilst lower values indicate random motion; “hurst_RS”: a metric of directional persistence. Values < 0.5 suggests non-persistent behaviour. A value of 0.5 indicates brownian motion. Values between >0.5 and 1.0 indicate persistent behaviour; “nongauss”: the extent of the non-Gaussian nature of the distribution of displacement of the cell within timelapse – value closer to 0 indicate Gaussian distribution; “rw_linearity”: linearity of a cell track minus linearity of a simulated random walk; “rw_netdist”: net distance travelled by a cell minus net distance of a simulated random walk; “rw_kurtosis”: kurtosis of a cell displacement minus kurtosis of a random walk for each sub-track; “avg_moving_speed” of a cell during a specified sub-track; “time_moving”: proportion of time spent moving by the cell during a sub-track; “autocorr”: similarity of a cell displacement series as a function of time lag between each displacement.

*In vitro* transendothelial migration (transwell) assay was performed using human umbilical vein endothelial cells (HUVECs; Lonza) grown at 37°C, 5% CO2 in EGM1 (Lonza) on 1% gelatin-coated flasks (Sigma), kept below 70% confluence and used up to passage 6. 8 μm porous cell culture membranes (BD Biosciences; 353093) were coated with 1.5% gelatin for 1 hour at 37°C, cross-linked with 2% glutaraldehyde (Sigma) for 15 minutes at RT, incubated with 70% ethanol for 1 hour at RT and washed 3 times with PBS before an overnight incubation in 2 mM glycine/PBS at 4°C. After 5 PBS washes, 2 × 10^5^ HUVECs were seeded on top and grown to confluence for at least 72 hours. hiMPs were then dissociated with TrypLE Select (Thermo Fisher Scientific) and labelled with 0.7 μM 6-carboxyfluorescin dictate (6-CFDA; Thermo Fisher Scientific) for 30 minutes at 37°C. Subsequently, the upper chamber was loaded with 3 × 10^4^ cells to be tested, resuspended in serum-free medium. The lower chamber was loaded with a chemoattractant composed of 50% fresh growth medium and 50% medium previously exposed for 24 hours to differentiated C2C12 myoblasts. After 8 hours, membranes were washed in PBS and fixed for 5 minutes in 4% PFA. The upper side of the membrane was scraped with a cotton bud to remove non-migrated cells. After an additional PBS wash, membranes were mounted on slides and cells migrated through the endothelial layer were quantified by counting the number of fluorescent cells on the lower side of the membrane using a Leica DMI6000B microscope. A minimum of 10 random 20X field/condition per experiment was quantified. Experiments were performed in duplicate on at least 3 separate occasions.

### Statistical analysis

Experiments were repeated at least three times prior to any statistical testing (“N” refers to independent experiments, “n” to data points). Quantification and statistical testing were performed using Microsoft Excel and GraphPad Prism 9 software. Statistical testing was based on Student’s t-test unless otherwise stated. Error bars on graphs represent standard error (average mean +/- SEM) unless otherwise stated. *P* values are specified in each figure on top of individual graphs.

## Supporting information

Supplemental file 1 (associated to figure 2E)

## ACKNOWLEDGEMENTS

The authors thank all members of the Tedesco laboratory, Yara Fadaili and Sara Benedetti for helpful feedback, UCL Genomics for support with initial RNAseq analyses, Shahragim Tajbakhsh and Hiroshi Sakai for providing mMuSCs from *Pax7-nGFP* mice. The MRC Centre for Neuromuscular Diseases Biobank, supported by the National Institute for Health Research (NIHR) Biomedical Research Centres at Great Ormond Street Hospital for Children NHS Foundation Trust and at University College London Hospitals NHS Foundation Trust and University College London, is acknowledged for providing human myoblasts. This work was funded by Muscular Dystrophy UK (RA4/3023/1, with Duchenne Children’s Trust / Duchenne UK and Duchenne Research Fund; 19GRO-PS48-0188; 17GRO-PS48-0093-1), the European Research Council (7591108 – HISTOID), the EU 7th Framework Program project no. 602423 (PluriMes) and the NIHR (the views expressed are those of the authors and not necessarily those of the National Health Service, the NIHR, or the Department of Health). Work in the Tedesco laboratory received also funding by the BBSRC, MRC, AFM-Telethon and Duchenne Parent Project.

## AUTHOR CONTRIBUTION

Conceptualisation, F.S.T. and G.F.; Methodology, G.F., S.W.C., L.A.M. and F.S.T.; Investigation, G.F., S.W.C., L.A.M., K.M., N.N., C.W., F.M. and F.S.T.; Writing, F.S.T., G.F. and S.W.C.; Funding acquisition and coordination, F.S.T.

## DECLARATION OF INTERESTS

The authors declare no conflict of interests.

## Notes

### Competing Interest Statement

The authors have declared no competing interest.

