## Supplemental information (figures and tables) for "DLL4 and PDGF-BB regulate migration of human iPSC-derived skeletal myogenic progenitors"

[1]Department of Cell and Developmental Biology, University College London, London, WC1E6DE, UK
[2]The Francis Crick Institute, 1 Midland Road, London, NW11AT, UK
[3]Sorbonne Université, Institut de Biologie Paris Seine FR3631, Plateforme de bioinformatique ARTbio, Paris, France.
[4]Centre for Stem Cell Systems, The University of Melbourne, Melbourne, Victoria, Australia.
[5]Dubowitz Neuromuscular Centre, UCL Great Ormond Street Institute of Child Health & Great Ormond Street Hospital for Children, London, UK
[6]National Institute for Health Research Great Ormond Street Hospital Biomedical Research Centre, London, UK
[#]Equally contributing authors
[§]Current address: Institute of Biomedical Engineering, University of Toronto, Toronto, M5S3E1, Canada
[Ø]Current address: University of Cambridge, Department of Physiology, Development and Neuroscience, Cambridge CB23DY, UK

- **Supplemental figures**
  - Figure S1
  - Figure S2
  - Figure S3
  - Figure S4
- **Supplemental tables**
  - Table S1
  - Table S2
  - Table S3
  - Table S4
- **Supplemental file** (online)

### Supplemental Figures

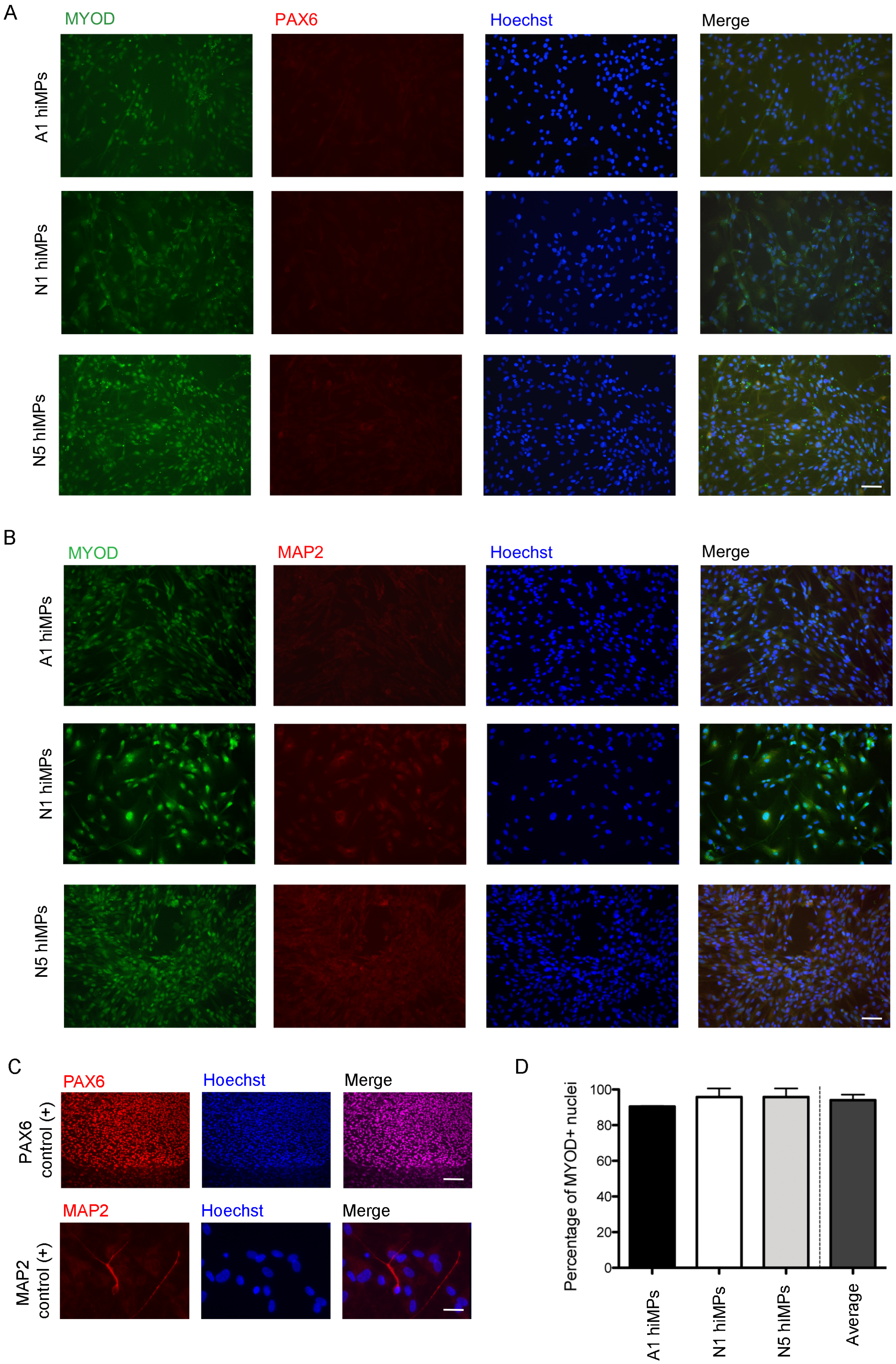


**Figure S1. Assessment of purity of hiMP populations.**

**(A)** Representative immunofluorescence analysis of MYOD (skeletal myogenic lineage marker, green) and PAX6 (early neuroectodermal lineage marker, red) immunoreactivity in three of the four different hiMP lines used in this study. **(B)** Immunofluorescence analysis of MYOD and MAP2 (late neuroectodermal / neuronal marker, red) in the same hiMPs shown in (A). **(C)** Positive controls for the PAX6 and MAP2 staining shown in (A, B); top panel: spontaneously differentiating hiPSCs; bottom panel: hiPSC-derived neurons. **(D)** Bar graph quantifying the percentages of MYOD-positive nuclei within three populations of hiMPs. Scale bars: (A, B) 75 µm; (C) top 100 µm; bottom 20 µm.

---

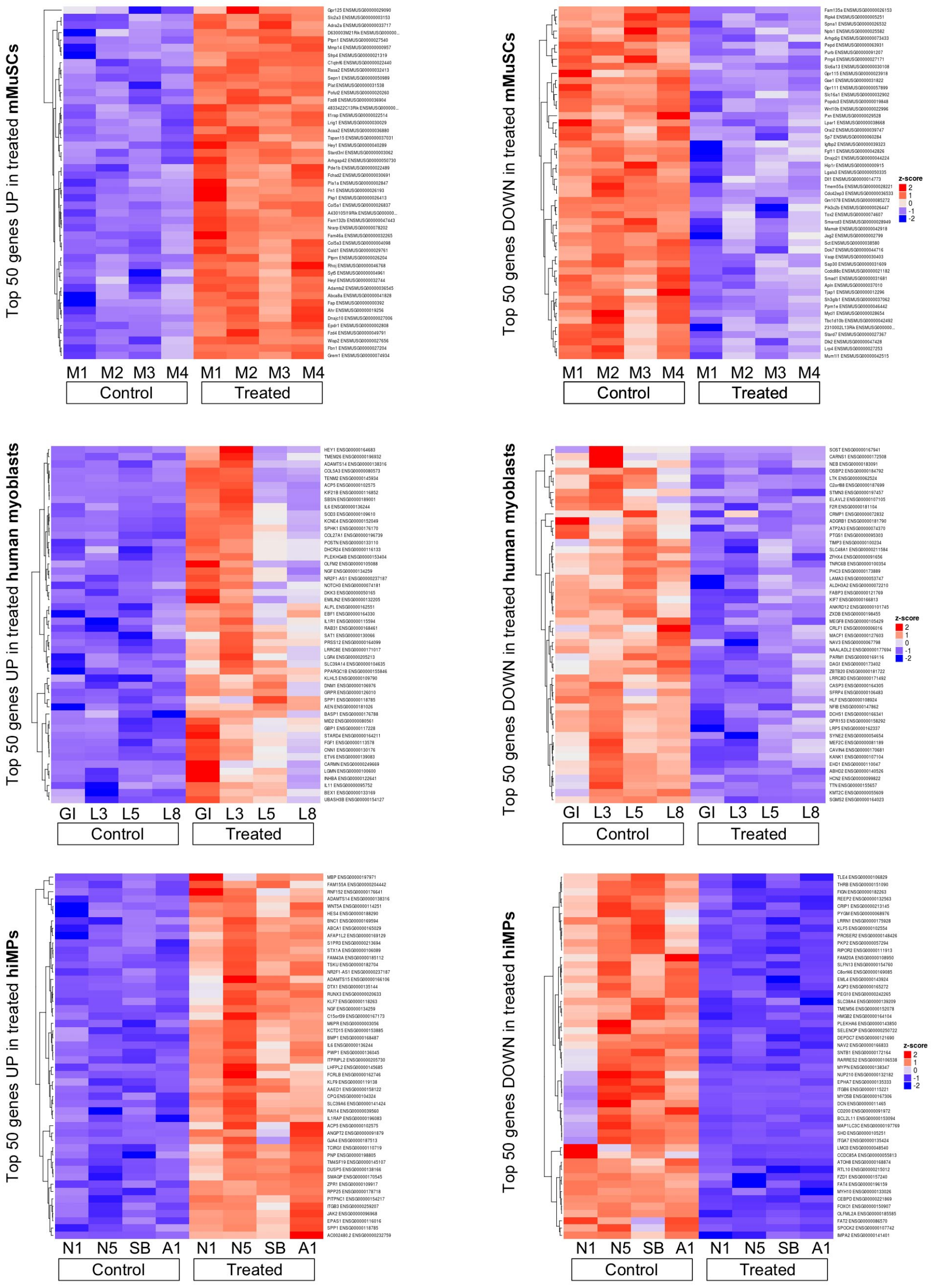


**Figure S2. Top 50 differentially regulated genes in mMuSC-derived myoblasts, human myoblasts and hiMPs.**

Heatmaps displaying 50 genes which exhibit either the greatest up- (left) or down-regulation (right) upon treatment with DLL4 & PDGF-BB in mMuSC (top), human myoblasts (centre) and hiMPs (bottom). Additional information (details on gene list) in Table S3.

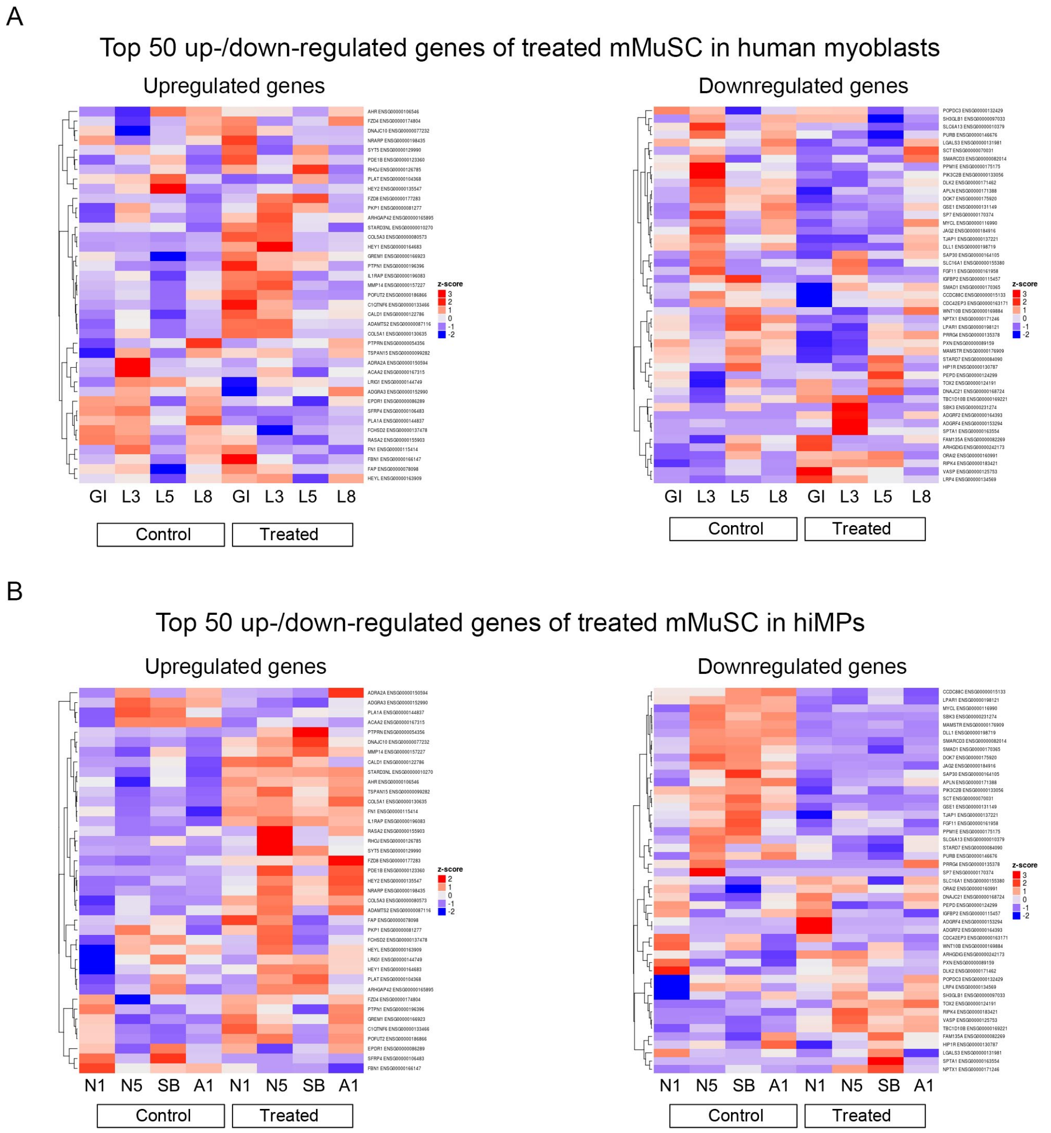


**Figure S3. Cross-comparison of top 50 differentially regulated genes of treated mMuSC-derived myoblasts in human myoblasts and hiMPs**

**(A)** Heatmaps of the top 50 up- (left) and down-regulated (right) genes of DLL4 & PDGF-BB-treated mMuSC-derived myoblasts in treated and untreated human myoblasts. **(B)** Heatmaps of the top 50 up- (left) and down-regulated (right) genes of DLL4 & PDGF-BB-treated mMuSC-derived myoblasts in treated and untreated hiMPs.

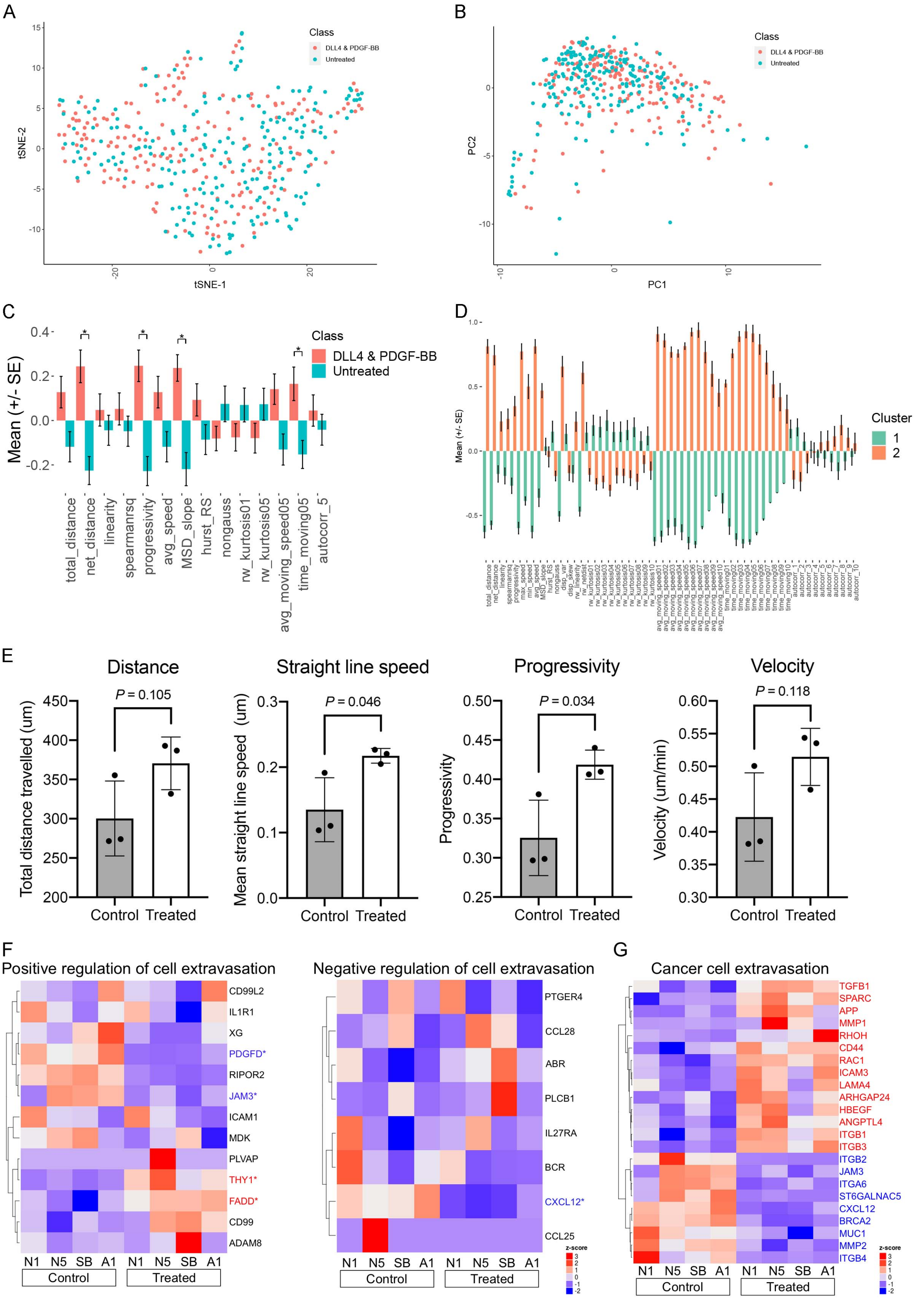


**Figure S4. Additional *in silico* and *in vitro* motility and migration analyses of treated and untreated hiMPs**

**(A)** t-SNE plot for visualisation of the motility state space occupied by DLL4 & PDGF-BB-treated cells (red) and untreated cells (light blue) (perplexity = 35). Cells obtained from 3 independent experimental replicates were analysed for each condition (n = 408). **(B)** Principal component analysis demonstrating the motility state space occupied by treated and untreated hiMPs within a linear space (n = 408). **(C)** Bar chart displaying normalised mean motility features for comparison of motility features between untreated and treated hiMPs (mean ±SEM). Statistical significance was based on a Bonferroni-corrected t-test. * *P* <0.05. **(D)** Bar chart comparing the extended normalised mean motility features for DLL4 & PDGF-BB-treated and untreated cells of the 2 clusters (mean ±SEM). **(E)** Bar graphs depict quantification of parameters obtained from single cell tracking analysed using TrackMate. Motility statistics were calculated for untreated (grey bars) and treated (white bars) hiMPs for 3 biological replicates (n = 3). *P* values within figure: *t*-test. **(F)** Heatmaps displaying genes that are involved in negative regulation of cellular extravasation (left; GO: 002692) and positive regulation of cellular extravasation (right; GO: 002693). **P* <0.05. **(G)** *P* value-adjusted hierarchical clustering heatmap showing a manually curated list of genes involved in enhanced trans-endothelial migration of cancer cells (*P* set al 0.05).

---

### Supplemental Tables

| | mMuSCs PC1 (45%) | mMuSCs PC2 (35%) | hMBs PC1(50%) | hMBs PC2 (25%) | hiMPs PC1 (63%) | hiMPs PC2 (17%) |
|---|---|---|---|---|---|---|
| 1 | Col15a1 | Il33 | COL5A3 | XIST | TTN | EPGN |
| 2 | Col6a2 | Actc1 | TRH | RPS4Y1 | MYBPH | CPA4 |
| 3 | Col1a1 | Flt1 | TTN | DDX3Y | SYNPO2 | SFRP1 |
| 4 | Col6a1 | Tek | SLC14A1 | USP9Y | TNNI1 | C3 |
| 5 | Grem1 | Jag1 | OLFM2 | NOS1 | MYH8 | UNC5B |
| 6 | Pdgfrb | Usp43 | POSTN | KDM5D | TNNT2 | SRGN |
| 7 | Heyl | Cyp2j6 | ASS1 | ZFY | CHRND | GDF6 |
| 8 | Bgn | Myog | CRISPLD2 | MYH3 | CHRNA1 | RGS4 |
| 9 | Adamts2 | Myo5b | ZNF469 | LINC00261 | KLHL41 | OXTR |
| 10 | Sfrp4 | Tnnt1 | ALDH1A1 | EIF1AY | SHD | IL1RL1 |
| 11 | Col6a3 | Tnnc2 | STMN2 | NLGN4Y | XIRP1 | MSC |
| 12 | Col5a3 | Atp2a1 | PTGIS | ERAP2 | ACTC1 | GUCY1A2 |
| 13 | Lrrc32 | C1qtnf3 | KLHL41 | ELN | RYR1 | HOXB9 |
| 14 | Cdh11 | Mylpf | MYH3 | HLA-A | ENO3 | INHBE |
| 15 | Nrarp | Grb10 | IGFBP5 | TXLNGY | CDH15 | ARRDC4 |
| 16 | Itgb3 | Nefm | INA | COL11A1 | UNC45B | TRIB3 |
| 17 | Thy1 | Sp7 | NOTCH3 | F13A1 | FNDC5 | CHRDL1 |
| 18 | Igfbp7 | Klhl41 | SERPINE2 | STMN2 | TNNC1 | DIO2 |
| 19 | Pkp1 | Smyd1 | KCNE4 | MMP1 | MYH3 | ZNF280D |
| 20 | Cyp1b1 | Lepr | JAG1 | EBF2 | SRL | PRKG1 |
| 21 | Mgp | Sema3d | LAMA3 | CLGN | DES | GPRC5C |
| 22 | Serping1 | Igf2 | MYBPH | TNNI1 | ACTA1 | NLGN1 |
| 23 | Ctgf | Meg3 | NEB | UTY | SMYD1 | NPY1R |
| 24 | Apbb1ip | Podxl | TENM2 | SFRP1 | MYH7 | DDIT3 |
| 25 | Fap | Synpo2l | NTSR1 | RARRES2 | VGLL2 | CCDC3 |
| 26 | Pcp4l1 | Rian | TNFRSF1B | TNNT2 | MYOZ2 | MSC-AS1 |
| 27 | Cldn4 | Zdbf2 | CD24 | PLXNA4 | MYOG | ADM2 |
| 28 | Igfbp2 | Myh3 | HSPB7 | KRT19 | LMO7 | DCC |
| 29 | Tcerg1l | Dchs1 | TNNT2 | IGFBP3 | MYOD1 | ENPP2 |
| 30 | Abcb1a | Nefl | MYLPF | COLEC12 | ACTN2 | TSPYL5 |
| 31 | Postn | Ldb3 | INHBA | ZNF185 | SFRP5 | UNC13A |
| 32 | S1pr1 | Fam84a | ADAMTS12 | PRKY | F13A1 | GATA6 |
| 33 | Tagln | Sfrp4 | NGFR | MYH7 | CKM | LAMC2 |
| 34 | Fn1 | Sct | KIF21B | SIM2 | NCAM1 | TYW3 |
| 35 | Trp53i11 | Gpnmb | MYOD1 | ANO1 | MYLPF | OLFM2 |
| 36 | Il6 | Aqp5 | SCG2 | IL17RD | STAC3 | SPON2 |
| 37 | Itgb5 | Myl1 | L1CAM | CECR1 | NEB | TUBB |
| 38 | Fam132b | Mstn | CKB | MYLPF | KLHL31 | KLF4 |
| 39 | Pde1b | H19 | F3 | TTTY15 | ITGA7 | BAALC |
| 40 | Scg2 | Actn2 | SFRP1 | ACTN2 | ERBB3 | DPP4 |
| 41 | Itga1 | Mybpc1 | COL4A1 | ACTA1 | GATM | LURAP1L |
| 42 | Cd248 | Btc | MYH7 | F2RL1 | MYL4 | TGM2 |
| 43 | Cd28 | Lmod3 | ADAM12 | IL13RA2 | MYPN | JRK |
| 44 | Slit2 | Myl4 | ALDH3A1 | HOXC10 | B3GALT2 | LGR4 |
| 45 | Pappa | Ppfia4 | ACTC1 | KIAA1462 | FGFR4 | MCTP2 |
| 46 | Gucy1a2 | Srl | CCDC141 | MYBPH | SHISA9 | GJB2 |
| 47 | Stc1 | Mylk4 | MYH8 | CASQ2 | NNAT | CXCL8 |
| 48 | Klf9 | Pdlim3 | COL5A1 | FLG | SORBS1 | CPE |
| 49 | Tnfaip2 | Nrep | ADAMTS2 | ANKRD1 | NPY | PCDH1 |
| 50 | Mrc2 | Acta1 | MYOG | SLIT2 | COL25A1 | TBX2 |

**Table S1.** Top 50 genes responsible for variations of PC1 and 2 in the principal component analysis shown in figure 2A.

| **50 top upregulated genes in mMuSCs** | **50 top up genes in human myoblasts** | **50 top upregulated genes in hiMPs** |
|---|---|---|
| Gpr125, ENSMUSG00000029090 | HEY1, ENSG00000164683 | MBP, ENSG00000197971 |
| Slc2a3, ENSMUSG00000003153 | TMEM26, ENSG00000196932 | FAM155A, ENSG00000204442 |
| Adra2a, ENSMUSG00000033717 | **ADAMTS14, ENSG00000138316** | RNF152, ENSG00000176641 |
| D630003M21Rik, ENSMUSG00000037813 | COL5A3, ENSG00000080573 | **ADAMTS14, ENSG00000138316** |
| Ptpn1, ENSMUSG00000027540 | TENM2, ENSG00000145934 | WNT5A, ENSG00000114251 |
| Mmp14, ENSMUSG00000000957 | ACP5, ENSG00000102575 | HES4, ENSG00000188290 |
| Sfrp4, ENSMUSG00000021319 | KIF21B, ENSG00000116852 | BNC1, ENSG00000169594 |
| C1qtnf6, ENSMUSG00000022440 | SBSN, ENSG00000189001 | ABCA1, ENSG00000165029 |
| Rasa2, ENSMUSG00000032413 | **IL6, ENSG00000136244** | AFAP1L2, ENSG00000169129 |
| Sepn1, ENSMUSG00000050989 | SOD3, ENSG00000109610 | S1PR3, ENSG00000213694 |
| Plat, ENSMUSG00000031538 | KCNE4, ENSG00000152049 | STX1A, ENSG00000106089 |
| Pofut2, ENSMUSG00000020260 | SPHK1, ENSG00000176170 | FAM43A, ENSG00000185112 |
| Fzd8, ENSMUSG00000036904 | COL27A1, ENSG00000196739 | TSKU, ENSG00000182704 |
| 4833422C13Rik, ENSMUSG00000074782 | POSTN, ENSG00000133110 | NR2F1-AS1, ENSG00000237187 |
| Il1rap, ENSMUSG00000022514 | DHCR24, ENSG00000116133 | **ADAMTS15, ENSG00000166106** |
| Lrig1, ENSMUSG00000030029 | PLEKHG4B, ENSG00000153404 | DTX1, ENSG00000135144 |
| Acaa2, ENSMUSG00000036880 | OLFM2, ENSG00000105088 | RUNX3, ENSG00000020633 |
| Tspan15, ENSMUSG00000037031 | **NGF, ENSG00000134259** | KLF7, ENSG00000118263 |
| Hey1, ENSMUSG00000040289 | **NR2F1-AS1, ENSG00000237187** | **NGF, ENSG00000134259** |
| Stard3nl, ENSMUSG00000003062 | NOTCH3, ENSG00000074181 | C15orf39, ENSG00000167173 |
| Arhgap42, ENSMUSG00000050730 | DKK3, ENSG00000050165 | M6PR, ENSG00000003056 |
| Pde1b, ENSMUSG00000022489 | EMILIN2, ENSG00000132205 | KCTD15, ENSG00000153885 |
| Fchsd2, ENSMUSG00000030691 | ALPL, ENSG00000162551 | BMP1, ENSG00000168487 |
| Pla1a, ENSMUSG00000002847 | EBF1, ENSG00000164330 | **IL6, ENSG00000136244** |
| Fn1, ENSMUSG00000026193 | IL1R1, ENSG00000115594 | PWP1, ENSG00000136045 |
| Pkp1, ENSMUSG00000026413 | RAB31, ENSG00000168461 | ITPRIPL2, ENSG00000205730 |
| Col5a1, ENSMUSG00000026837 | SAT1, ENSG00000130066 | LHFPL2, ENSG00000145685 |
| A430105I19Rik, ENSMUSG00000045838 | PRSS12, ENSG00000164099 | FCRLB, ENSG00000162746 |
| Fam132b, ENSMUSG00000047443 | LRRC8E, ENSG00000171017 | KLF9, ENSG00000119138 |
| Nrarp, ENSMUSG00000078202 | LGR4, ENSG00000205213 | AAED1, ENSG00000158122 |
| Fam46a, ENSMUSG00000032265 | SLC39A14, ENSG00000104635 | CPQ, ENSG00000104324 |
| Col5a3, ENSMUSG00000004098 | PPARGC1B, ENSG00000155846 | SLC39A6, ENSG00000141424 |
| Cald1, ENSMUSG00000029761 | KLHL5, ENSG00000109790 | RAI14, ENSG00000039560 |
| Ptprn, ENSMUSG00000026204 | DNM1, ENSG00000106976 | IL1RAP, ENSG00000196083 |
| Rhoj, ENSMUSG00000046768 | GRPR, ENSG00000126010 | ACP5, ENSG00000102575 |
| Syt5, ENSMUSG00000004961 | **SPP1, ENSG00000118785** | ANGPT2, ENSG00000091879 |
| Heyl, ENSMUSG00000032744 | AEN, ENSG00000181026 | GJA4, ENSG00000187513 |
| Adamts2, ENSMUSG00000036545 | BASP1, ENSG00000176788 | TCIRG1, ENSG00000110719 |
| Abca8a, ENSMUSG00000041828 | MID2, ENSG00000080561 | PNP, ENSG00000198805 |
| Fap, ENSMUSG00000000392 | GBP1, ENSG00000117228 | TM4SF19, ENSG00000145107 |
| Ahr, ENSMUSG00000019256 | STARD4, ENSG00000164211 | DUSP5, ENSG00000138166 |
| Dnajc10, ENSMUSG00000027006 | FGF1, ENSG00000113578 | SMAGP, ENSG00000170545 |
| Epdr1, ENSMUSG00000002808 | CNN1, ENSG00000130176 | ZPR1, ENSG00000109917 |
| Fzd4, ENSMUSG00000049791 | ETV6, ENSG00000139083 | RPP25, ENSG00000178718 |
| Wisp2, ENSMUSG00000027656 | CARMN, ENSG00000249669 | PITPNC1, ENSG00000154217 |
| Fbn1, ENSMUSG00000027204 | LGMN, ENSG00000100600 | ITGB3, ENSG00000259207 |
| Grem1, ENSMUSG00000074934 | INHBA, ENSG00000122641 | JAK2, ENSG00000096968 |
| *Cfh ENSMUSG00000026365 | IL11, ENSG00000095752 | EPAS1, ENSG00000116016 |
| *AW011738, ENSMUSG00000078349 | BEX1, ENSG00000133169 | **SPP1, ENSG00000118785** |
| *Hey2, ENSMUSG00000019789 | UBASH3B, ENSG00000154127 | AC002480.2, ENSG00000232759 |

| **50 top downregulated genes in mMuSCs** | **50 top downregulated genes in human myoblasts** | **50 top downregulated genes in hiMPs** |
|---|---|---|
| Fam135a, ENSMUSG00000026153 | SOST, ENSG00000167941 | TLE4, ENSG00000106829,, |
| Ripk4, ENSMUSG00000005251 | CARNS1, ENSG00000172508 | THRB, ENSG00000151090 |
| Spna1, ENSMUSG00000026532 | NEB, ENSG00000183091 | FIGN, ENSG00000182263 |
| Nptx1, ENSMUSG00000025582 | OSBP2, ENSG00000184792 | REEP2, ENSG00000132563 |
| Arhgdig, ENSMUSG00000073433 | LTK, ENSG00000062524 | CRIP1, ENSG00000213145 |
| Pepd, ENSMUSG00000063931 | C2orf88, ENSG00000187699 | PYGM, ENSG00000068976 |
| Purb, ENSMUSG00000091207 | STMN3, ENSG00000197457 | LRRN1, ENSG00000175928 |
| Prrg4, ENSMUSG00000027171 | ELAVL2, ENSG00000107105 | KLF5, ENSG00000102554 |
| Slc6a13, ENSMUSG00000030108 | F2R, ENSG00000181104 | PROSER2, ENSG00000148426 |
| Gpr115, ENSMUSG00000023918 | CRMP1, ENSG00000072832 | PKP2, ENSG00000057294 |
| Gse1, ENSMUSG00000031822 | ADGRB1, ENSG00000181790 | RIPOR2, ENSG00000111913 |
| Gpr111, ENSMUSG00000057899 | ATP2A3, ENSG00000074370 | FAM20A, ENSG00000108950 |
| Slc16a1, ENSMUSG00000032902 | PTGS1, ENSG00000095303 | SLFN13, ENSG00000154760 |
| Popdc3, ENSMUSG00000019848 | TIMP3, ENSG00000100234 | C8orf46, ENSG00000169085 |
| Wnt10b, ENSMUSG00000022996 | SLC48A1, ENSG00000211584 | EML4, ENSG00000143924 |
| Pxn, ENSMUSG00000029528 | ZFHX4, ENSG00000091656 | AQP3, ENSG00000165272 |
| Lpar1, ENSMUSG00000038668 | TNRC6B, ENSG00000100354 | PEG10, ENSG00000242265 |
| Orai2, ENSMUSG00000039747 | PHC3, ENSG00000173889 | SLC38A4, ENSG00000139209 |
| Sp7, ENSMUSG00000060284 | LAMA3, ENSG00000053747 | TMEM56, ENSG00000152078 |
| Igfbp2, ENSMUSG00000039323 | ALDH3A2, ENSG00000072210 | HMGB2, ENSG00000164104 |
| Fgf11, ENSMUSG00000042826 | FABP3, ENSG00000121769 | PLEKHA6, ENSG00000143850 |
| Dnajc21, ENSMUSG00000044224 | KIF7, ENSG00000166813 | SELENOP, ENSG00000250722 |
| Hip1r, ENSMUSG00000000915 | ANKRD12, ENSG00000101745 | DEPDC7, ENSG00000121690 |
| Lgals3, ENSMUSG00000050335 | ZXDB, ENSG00000198455 | NAV2, ENSG00000166833 |
| Dll1, ENSMUSG00000014773 | MEGF8, ENSG00000105429 | SNTB1, ENSG00000172164 |
| Tmem55a, ENSMUSG00000028221 | CRLF1, ENSG00000006016 | RARRES2, ENSG00000106538 |
| Cdc42ep3, ENSMUSG00000036533 | MACF1, ENSG00000127603 | MYPN, ENSG00000138347 |
| Gm1078, ENSMUSG00000085272 | NAV3, ENSG00000067798 | NUP210, ENSG00000132182 |
| Pik3c2b, ENSMUSG00000026447 | NAALADL2, ENSG00000177694 | EPHA7, ENSG00000135333 |
| Tox2, ENSMUSG00000074607 | PARM1, ENSG00000169116 | ITGB6, ENSG00000115221 |
| Smarcd3, ENSMUSG00000028949 | DAG1, ENSG00000173402 | MYO5B, ENSG00000167306 |
| Mamstr, ENSMUSG00000042918 | ZBTB20, ENSG00000181722 | DCN, ENSG00000011465 |
| Jag2, ENSMUSG00000002799 | LRRC8D, ENSG00000171492 | CD200, ENSG00000091972 |
| Sct, ENSMUSG00000038580 | CASP3, ENSG00000164305 | BCL2L11, ENSG00000153094 |
| Dok7, ENSMUSG00000044716 | SFRP4, ENSG00000106483 | MAP1LC3C, ENSG00000197769 |
| Vasp, ENSMUSG00000030403 | HLF, ENSG00000108924 | SHD, ENSG00000105251 |
| Sap30, ENSMUSG00000031609 | NFIB, ENSG00000147862 | ITGA7, ENSG00000135424 |
| Ccdc88c, ENSMUSG00000021182 | DCHS1, ENSG00000166341 | LMO3, ENSG00000048540 |
| Smad1, ENSMUSG00000031681 | GPR153, ENSG00000158292 | CCDC85A, ENSG00000055813 |
| Apln, ENSMUSG00000037010 | LRP5, ENSG00000162337 | ATOH8, ENSG00000168874 |
| Tjap1, ENSMUSG00000012296 | SYNE2, ENSG00000054654 | RTL10, ENSG00000215012 |
| Sh3glb1, ENSMUSG00000037062 | MEF2C, ENSG00000081189 | FZD1, ENSG00000157240 |
| Ppm1e, ENSMUSG00000046442 | CAVIN4, ENSG00000170681 | FAT4, ENSG00000196159 |
| Mycl1, ENSMUSG00000028654 | KANK1, ENSG00000107104 | MYH10, ENSG00000133026 |
| Tbc1d10b, ENSMUSG00000042492 | EHD1, ENSG00000110047 | CEBPD, ENSG00000221869 |
| 2310002L13Rik, ENSMUSG00000024512 | ABHD2, ENSG00000140526 | FOXO1, ENSG00000150907 |
| Stard7, ENSMUSG00000027367 | HCN2, ENSG00000099822 | OLFML2A, ENSG00000185585 |
| Dlk2, ENSMUSG00000047428 | TTN, ENSG00000155657 | FAT2, ENSG00000086570 |
| Lrp4, ENSMUSG00000027253 | KMT2C, ENSG00000055609 | SPOCK2, ENSG00000107742 |
| Mum1l1, ENSMUSG00000042515 | SGMS2, ENSG00000164023 | IMPA2, ENSG00000141401 |

**Table S2.** Full list of genes of heatmaps shown in figure S2 displaying 50 genes which exhibit either the greatest up- or down-regulation upon treatment with DLL4 & PDGF-BB in mMuSC (left), human myoblasts (centre) and hiMPs (right). *Genes not shown in heatmap due to N/A rows resulting from Stemformatics analysis. Bold font: common genes in human lists.

| **Ranked genes** | | | | | | | |
|---|---|---|---|---|---|---|---|
| **Top 50 upregulated genes of treated mMuSC in human myoblasts** | | **Top 50 downregulated genes of treated mMuSC in human myoblasts** | | **Top 50 upregulated genes of treated mMuSC in hiMPs** | | **Top 50 downregulated genes of treated mMuSC in hiMPs** | |
| Gene | Probe | Gene | Probe | Gene | Probe | Gene | Probe |
| AHR | ENSG00000106546 | POPDC3 | ENSG00000132429 | ADRA2A | ENSG00000150594 | CCDC88C | ENSG00000015133 |
| FZD4 | ENSG00000174804 | SH3GLB1 | ENSG00000097033 | ADGRA3 | ENSG00000152990 | LPAR1 | ENSG00000198121 |
| DNAJC10 | ENSG00000077232 | SLC6A13 | ENSG00000010379 | PLA1A | ENSG00000144837 | MYCL | ENSG00000116990 |
| NRARP | ENSG00000198435 | PURB | ENSG00000146676 | ACAA2 | ENSG00000167315 | SBK3 | ENSG00000231274 |
| SYT5 | ENSG00000129990 | LGALS3 | ENSG00000131981 | PTPRN | ENSG00000054356 | MAMSTR | ENSG00000176909 |
| PDE1B | ENSG00000123360 | SCT | ENSG00000070031 | DNAJC10 | ENSG00000077232 | DLL1 | ENSG00000198719 |
| RHOJ | ENSG00000126785 | SMARCD3 | ENSG00000082014 | MMP14 | ENSG00000157227 | SMARCD3 | ENSG00000082014 |
| PLAT | ENSG00000104368 | PPM1E | ENSG00000175175 | CALD1 | ENSG00000122786 | SMAD1 | ENSG00000170365 |
| HEY2 | ENSG00000135547 | PIK3C2B | ENSG00000133056 | STARD3NL | ENSG00000010270 | DOK7 | ENSG00000175920 |
| FZD8 | ENSG00000177283 | DLK2 | ENSG00000171462 | AHR | ENSG00000106546 | JAG2 | ENSG00000184916 |
| PKP1 | ENSG00000081277 | APLN | ENSG00000171388 | TSPAN15 | ENSG00000099282 | SAP30 | ENSG00000164105 |
| ARHGAP42 | ENSG00000165895 | DOK7 | ENSG00000175920 | COL5A1 | ENSG00000130635 | APLN | ENSG00000171388 |
| STARD3NL | ENSG00000010270 | GSE1 | ENSG00000131149 | FN1 | ENSG00000115414 | PIK3C2B | ENSG00000133056 |
| COL5A3 | ENSG00000080573 | SP7 | ENSG00000170374 | IL1RAP | ENSG00000196083 | SCT | ENSG00000070031 |
| HEY1 | ENSG00000164683 | MYCL | ENSG00000116990 | RASA2 | ENSG00000155903 | GSE1 | ENSG00000131149 |
| GREM1 | ENSG00000166923 | JAG2 | ENSG00000184916 | RHOJ | ENSG00000126785 | TJAP1 | ENSG00000137221 |
| PTPN1 | ENSG00000196396 | TJAP1 | ENSG00000137221 | SYT5 | ENSG00000129990 | FGF11 | ENSG00000161958 |
| IL1RAP | ENSG00000196083 | DLL1 | ENSG00000198719 | FZD8 | ENSG00000177283 | PPM1E | ENSG00000175175 |
| MMP14 | ENSG00000157227 | SAP30 | ENSG00000164105 | PDE1B | ENSG00000123360 | SLC6A13 | ENSG00000010379 |
| POFUT2 | ENSG00000186866 | SLC16A1 | ENSG00000155380 | HEY2 | ENSG00000135547 | STARD7 | ENSG00000084090 |
| C1QTNF6 | ENSG00000133466 | FGF11 | ENSG00000161958 | NRARP | ENSG00000198435 | PURB | ENSG00000146676 |
| CALD1 | ENSG00000122786 | IGFBP2 | ENSG00000115457 | COL5A3 | ENSG00000080573 | PRRG4 | ENSG00000135378 |
| ADAMTS2 | ENSG00000087116 | SMAD1 | ENSG00000170365 | ADAMTS2 | ENSG00000087116 | SP7 | ENSG00000170374 |
| COL5A1 | ENSG00000130635 | CCDC88C | ENSG00000015133 | FAP | ENSG00000078098 | SLC16A1 | ENSG00000155380 |
| PTPRN | ENSG00000054356 | CDC42EP3 | ENSG00000163171 | PKP1 | ENSG00000081277 | ORAI2 | ENSG00000160991 |
| TSPAN15 | ENSG00000099282 | WNT10B | ENSG00000169884 | FCHSD2 | ENSG00000137478 | DNAJC21 | ENSG00000168724 |
| ADRA2A | ENSG00000150594 | NPTX1 | ENSG00000171246 | HEYL | ENSG00000163909 | PEPD | ENSG00000124299 |
| ACAA2 | ENSG00000167315 | LPAR1 | ENSG00000198121 | LRIG1 | ENSG00000144749 | IGFBP2 | ENSG00000115457 |
| LRIG1 | ENSG00000144749 | PRRG4 | ENSG00000135378 | HEY1 | ENSG00000164683 | ADGRF4 | ENSG00000153294 |
| ADGRA3 | ENSG00000152990 | PXN | ENSG00000089159 | PLAT | ENSG00000104368 | ADGRF2 | ENSG00000164393 |
| EPDR1 | ENSG00000086289 | MAMSTR | ENSG00000176909 | ARHGAP42 | ENSG00000165895 | CDC42EP3 | ENSG00000163171 |
| SFRP4 | ENSG00000106483 | STARD7 | ENSG00000084090 | FZD4 | ENSG00000174804 | WNT10B | ENSG00000169884 |
| PLA1A | ENSG00000144837 | HIP1R | ENSG00000130787 | PTPN1 | ENSG00000196396 | ARHGDIG | ENSG00000242173 |
| FCHSD2 | ENSG00000137478 | PEPD | ENSG00000124299 | GREM1 | ENSG00000166923 | PXN | ENSG00000089159 |
| RASA2 | ENSG00000155903 | TOX2 | ENSG00000124191 | C1QTNF6 | ENSG00000133466 | DLK2 | ENSG00000171462 |
| FN1 | ENSG00000115414 | DNAJC21 | ENSG00000168724 | POFUT2 | ENSG00000186866 | POPDC3 | ENSG00000132429 |
| FBN1 | ENSG00000166147 | TBC1D10B | ENSG00000169221 | EPDR1 | ENSG00000086289 | LRP4 | ENSG00000134569 |
| FAP | ENSG00000078098 | SBK3 | ENSG00000231274 | SFRP4 | ENSG00000106483 | SH3GLB1 | ENSG00000097033 |
| HEYL | ENSG00000163909 | ADGRF2 | ENSG00000164393 | FBN1 | ENSG00000166147 | TOX2 | ENSG00000124191 |
| Fam132b | * | ADGRF4 | ENSG00000153294 | Fam132b | * | RIPK4 | ENSG00000183421 |
| 4833422C13Rik | * | SPTA1 | ENSG00000163554 | 4833422C13Rik | * | VASP | ENSG00000125753 |
| Wisp2 | * | FAM135A | ENSG00000082269 | Wisp2 | * | TBC1D10B | ENSG00000169221 |
| A430105I19Rik | * | ARHGDIG | ENSG00000242173 | A430105I19Rik | * | FAM135A | ENSG00000082269 |
| Slc2a3 | * | ORAI2 | ENSG00000160991 | Slc2a3 | * | HIP1R | ENSG00000130787 |
| Abca8a | * | RIPK4 | ENSG00000183421 | Abca8a | * | LGALS3 | ENSG00000131981 |
| Cfh | * | VASP | ENSG00000125753 | Cfh | * | SPTA1 | ENSG00000163554 |
| Fam46a | * | LRP4 | ENSG00000134569 | Fam46a | * | NPTX1 | ENSG00000171246 |
| Sepn1 | * | Tmem55a | * | Sepn1 | * | Tmem55a | * |
| AW011738 | * | Mum1l1 | * | AW011738 | * | Mum1l1 | * |
| D630003M21Rik | * | Dynap | * | D630003M21Rik | * | Dynap | * |

**Table S3.** Table of ranked genes supplementing heatmaps presented in Figure S3: “Top 50 upregulated genes of treated mMuSC-derived myoblasts in human myoblasts” (Figure S3A) (left); “Top 50 downregulated genes of treated mMuSC-derived myoblasts in human myoblasts” (Figure S3A) (right); “Top 50 upregulated genes of treated mMuSC-derived myoblasts in hiMPs” (Figure S3B) (left); “Top 50 downregulated genes of treated mMuSC-derived myoblasts in hiMPs” (Figure S3B) (right). *no human orthologue found.

| Ranked genes | | | | | |
|---|---|---|---|---|---|
| **Regulation of cell morphology** | | **Proliferation of stem/myogenic Cells** | | **Leukocyte trans-endothelial migration** | |
| Gene | Probe | Gene | Probe | Gene | Probe |
| MYO10 | ENSG00000145555 | MMP9 | ENSG00000100985 | TXK | ENSG00000074966 |
| VEGFA | ENSG00000112715 | PTGIR | ENSG00000160013 | ACTN3 | ENSG00000248746 |
| RHOQ | ENSG00000119729 | VEGFA | ENSG00000112715 | VAV3 | ENSG00000134215 |
| FN1 | ENSG00000115414 | PDGFRB | ENSG00000113721 | CLDN5 | ENSG00000184113 |
| KIT | ENSG00000157404 | NGF | ENSG00000134259 | ITGB2 | ENSG00000160255 |
| RAC3 | ENSG00000169750 | IRAK1 | ENSG00000184216 | RASSF5 | ENSG00000266094 |
| WIPF1 | ENSG00000115935 | TGFB1 | ENSG00000105329 | ACTN2 | ENSG00000077522 |
| SH3KBP1 | ENSG00000147010 | C3AR1 | ENSG00000171860 | JAM2 | ENSG00000154721 |
| RHOC | ENSG00000155366 | KITLG | ENSG00000049130 | PIK3R1 | ENSG00000145675 |
| HEXB | ENSG00000049860 | GNAI3 | ENSG00000065135 | JAM3 | ENSG00000166086 |
| CDC42EP4 | ENSG00000179604 | SIRT6 | ENSG00000077463 | MYLPF | ENSG00000180209 |
| RAC1 | ENSG00000136238 | JAK2 | ENSG00000096968 | CTNNA3 | ENSG00000183230 |
| MSN | ENSG00000147065 | ITGB3 | ENSG00000259207 | GNAI1 | ENSG00000127955 |
| PLXND1 | ENSG00000004399 | SNAI2 | ENSG00000019549 | CXCL12 | ENSG00000107562 |
| IL6 | ENSG00000136244 | IL6 | ENSG00000136244 | MYL2 | ENSG00000111245 |
| FMNL3 | ENSG00000161791 | NOTCH3 | ENSG00000074181 | MMP9 | ENSG00000100985 |
| MYH9 | ENSG00000100345 | MYC | ENSG00000136997 | PLCG2 | ENSG00000197943 |
| KIF3A | ENSG00000131437 | NOS3 | ENSG00000164867 | CLDN7 | ENSG00000181885 |
| FBLIM1 | ENSG00000162458 | ILK | ENSG00000166333 | ESAM | ENSG00000149564 |
| CDC42EP1 | ENSG00000128283 | TRIB1 | ENSG00000173334 | ARHGAP35 | ENSG00000160007 |
| DLC1 | ENSG00000164741 | IL12A | ENSG00000168811 | RHOH | ENSG00000168421 |
| ARHGAP35 | ENSG00000160007 | HBEGF | ENSG00000113070 | ACTN1 | ENSG00000072110 |
| ARAP3 | ENSG00000120318 | CAV1 | ENSG00000105974 | ITGB1 | ENSG00000150093 |
| RHOG | ENSG00000177105 | CNN1 | ENSG00000130176 | VASP | ENSG00000125753 |
| RHOD | ENSG00000173156 | FGF9 | ENSG00000102678 | THY1 | ENSG00000154096 |
| SEMA4D | ENSG00000187764 | BMPR1A | ENSG00000107779 | RAP1B | ENSG00000127314 |
| LPAR1 | ENSG00000198121 | HMGB2 | ENSG00000164104 | RAC1 | ENSG00000136238 |
| MYH10 | ENSG00000133026 | BMP4 | ENSG00000125378 | GNAI3 | ENSG00000065135 |
| RHOBTB3 | ENSG00000164292 | RBPJ | ENSG00000168214 | MSN | ENSG00000147065 |
| PHIP | ENSG00000146247 | CTNNBIP1 | ENSG00000178585 | | |
| S100A13 | ENSG00000189171 | SOX15 | ENSG00000129194 | | |
| PALMD | ENSG00000099260 | MYOD1 | ENSG00000129152 | | |
| ITGA7 | ENSG00000135424 | MAP3K5 | ENSG00000197442 | | |
| PALM2 | ENSG00000243444 | PIK3R1 | ENSG00000145675 | | |
| EPB41L3 | ENSG00000082397 | MYOG | ENSG00000122180 | | |
| WIPF3 | ENSG00000122574 | KLHL41 | ENSG00000239474 | | |
| SEMA3E | ENSG00000170381 | SMARCD3 | ENSG00000082014 | | |
| KDR | ENSG00000128052 | MAGI1 | ENSG00000151276 | | |
| PLXNB1 | ENSG00000164050 | CAMK2D | ENSG00000145349 | | |
| | | MEF2C | ENSG00000081189 | | |
| | | MEGF10 | ENSG00000145794 | | |
| | | PPARGC1A | ENSG00000109819 | | |
| | | PDE1A | ENSG00000115252 | | |
| | | ANGPT1 | ENSG00000154188 | | |
| | | MMP2 | ENSG00000087245 | | |
| | | RGS5 | ENSG00000143248 | | |
| | | IL18 | ENSG00000150782 | | |
| | | PDGFD | ENSG00000170962 | | |
| | | AKR1B1 | ENSG00000085662 | | |
| | | MNAT1 | ENSG00000020426 | | |
| | | SKP2 | ENSG00000145604 | | |
| | | EGR1 | ENSG00000120738 | | |
| | | TGM2 | ENSG00000198959 | | |
| | | DNMT1 | ENSG00000130816 | | |
| | | ASPM | ENSG00000066279 | | |
| | | ORC1 | ENSG00000085840 | | |

**Table S4.** Table of ranked genes accompanying heatmaps “Regulation of cell morphology”, “Proliferation of stem/myogenic cells” and “Leukocyte trans-endothelial migration” (Figures 3A, 3D and 4G, respectively).

---

**Supplemental File 1.** Spreadsheet containing the three full lists of the enrichment analyses shown in figure 2E. *P* value set at 0.05 (available online).

**END OF DOCUMENT**
